# Amyloid β-peptides interfere with mitochondrial preprotein import competence by a co-aggregation process

**DOI:** 10.1101/050617

**Authors:** Giovanna Cenini, Cornelia Rüb, Michael Bruderek, Wolfgang Voos

## Abstract

Aβ peptides play a central role in the etiology of Alzheimer disease (AD) by exerting cellular toxicity correlated with aggregate formation. Experimental evidences showed an intraneuronal accumulation of Aβ peptides and an interference with mitochondrial functions. Nevertheless, the relevance of intracellular Aβ peptides in the pathophysiology of AD remained controversial. Here, we found that the two major species of Aβ peptides, in particular Aβ42, exhibited a strong inhibitory effect on the preprotein import reactions essential for mitochondrial biogenesis. However, Aβ peptides interacted only weakly with mitochondria and did not affect the inner membrane potential or the structure of the preprotein translocase complexes. Aβ peptides significantly decreased the import competence of mitochondrial precursor proteins through a extra-mitochondrial co-aggregation mechanism. Co-aggregation and import inhibition were significantly stronger in case of the longer peptide Aβ42, correlating with its importance in AD pathology. Our results demonstrate that a direct interference of aggregation-prone Aβ peptides with mitochondrial protein biogenesis represents a crucial aspect of the pathobiochemical mechanisms contributing to cellular damage in AD.

**List of Abbreviations**
TMRE: tetramethylrhodamine ethyl ester
TCA: trichloroacetic acid
MPP: mitochondrial processing peptidase
BN-PAGE: blue-native polyacrylamide gel electrophoresis
SDS-PAGE: sodiumdodecylsulfate polyacrylamide gel electrophoresis
TOM: translocase of the outer membrane
TIM: translocase of the inner membrane
OMM: outer mitochondrial membrane
IMM: inner mitochondrial membrane
IMS: intermembrane space
DHFR: dihydrofolate reductase
AA: amino acid
Δψ_mt_: electric potential across the inner mitochondrial membrane

## Introduction

Beta-amyloid (Aβ) peptides have been associated with severe human pathological conditions like Alzheimer disease (AD) (Murphy and LeVine, 2010), Down syndrome (Head and Lott, 2004) and cerebral amyloid angiopathy (Weller *et al.*, 2000), all characterized by accumulation and deposition of Aβ peptides in the central nervous system. Due to the diversity of pathological aspects connected with a severe neurodegenerative disease like AD, the biochemical mechanisms resulting in neuronal cell death and the correlation with the accumulation of Aβ peptides are not completely clear (Musiek and Holtzman, 2015).

Aβ peptides derive from a proteolytic process mediated by β- and γ-secretases on the type 1 trans-membrane precursor called amyloid precursor protein (APP). The most common forms in AD are constituted of 40 (Aβ40) and 42 (Aβ42) amino acids (Zhang *et al.*, 2011). Mutations, environmental factors as well as aging could induce changes in the equilibrium between Aβ peptide production and removal (Mawuenyega *et al.*, 2010) as well as an imbalance between amyloidogenic and non-amyloidogenic pathways (Agostinho *et al.*, 2015). This causes an increase of Aβ peptide concentrations promoting aggregation and deposition as senile plaques in brain parenchyma. Kinetic and structural studies about Aβ aggregation *in vitro* have reported that unstructured Aβ monomers have an intrinsic tendency to self-assemble spontaneously by a nucleation-polymerization mechanism into higher-order oligomeric, protofibrillar and fibrillar states (Thal *et al.*, 2015). The aggregation process is enhanced by high peptide concentrations, presence of nucleation seeds, altered pH, ionic strength, or temperature (Stine *et al.*, 2003). Furthermore, a large variety of post-translation modifications of the Aβ sequence influence the aggregation propensity (Kummer and Heneka, 2014; Thal *et al.*, 2015). As Aβ42 oligomers represent the most toxic amyloidogenic peptide species, the main component of AD senile plaques, and the first to deposit during the senile plaques formation, they play a key pathophysiological role in the development of AD (Haass and Selkoe, 2007). Interestingly, although Aβ42 has only small structural differences compared to the other Aβ peptides, it displays distinct clinical, biological and biophysical behaviors (Jarrett *et al.*, 1993; Bitan *et al.*, 2003).

The “amyloid cascade hypothesis” represents the major theory to explain the etiology and pathology of AD (Hardy and Selkoe, 2002; Musiek and Holtzman, 2015). This hypothesis, strongly supported by genetic studies on familial AD cases (Hardy and Higgins, 1992), proposed that an aggregation of Aβ peptides is responsible for the initiation of a multistep pathological cascade eventually resulting in neuronal death. A growing body of evidence also suggested the prominent contribution of an intracellular accumulation of Aβ peptides as a trigger of neurodegeneration and AD pathology on the cellular level (Wirths *et al.*, 2004; Gouras *et al.*, 2010; Wirths and Bayer, 2012). Intracellular pools of Aβ peptides may stem from an intracellular production, by a reuptake of secreted peptide molecules or both. Eventually accumulating also in the cytosol, it is likely that intracellular Aβ peptides interact with membranes or other cellular components and induce structural changes of sub-cellular compartments (LaFerla *et al.*, 2007).

Mitochondrial dysfunction is now consensually accepted as a general pathological feature in AD patients (Mattson *et al.*, 2008; Piaceri *et al.*, 2012; Selfridge *et al.*, 2013). In line with this, a modification of the amyloid cascade hypothesis was postulated that supports the correlation between mitochondrial dysfunction with AD. Named “mitochondrial cascade hypothesis”, it considers how individual mitochondrial dysfunctions, accumulating in aging cells, could influence Aβ peptide homeostasis, aggregation and consequently the chronology of AD (Swerdlow *et al.*, 2014). However, it is still disputed if mitochondrial dysfunctions are early casual events or a consequence of other pathological events in AD patients. Evidences exist that indicate an accumulation of Aβ peptides in mitochondria, interactions with protein components of the mitochondrial matrix, and perturbations of mitochondrial functions (Lustbader *et al.*, 2004; Hansson Petersen *et al.*, 2008; Mossmann *et al.*, 2014; Kaminsky *et al.*, 2015). Nevertheless, the molecular mechanisms underlying a mitochondrial accumulation and the claimed effects of Aβ peptides on mitochondria need a critical analysis and clarification. For this reason, we performed a comprehensive biochemical analysis of the interaction between the two Aβ peptides species relevant to AD (Aβ40 and Aβ42) with human mitochondria. One of the major cellular processes responsible for maintaining mitochondrial functions is the import of nuclear-encoded mitochondrial precursor proteins from the cytosol (Chacinska *et al.*, 2009). We utilized an established import assay based on isolated intact mitochondria (Ryan *et al.*, 2001) to check if and how Aβ peptides directly interfere with the mitochondrial protein import reaction. Taken together, our results show a strong and direct inhibitory effect of Aβ peptides on mitochondrial protein biogenesis. This inhibition is not caused by a damaging influence of Aβ peptides on mitochondrial functions, but is correlated to a co-aggregation phenomenon between Aβ peptides and precursor proteins that severely restricts their import competence.

## Results

### Aβ peptides interfere with the import of mitochondrial precursor proteins

The import of precursor proteins, synthesized at cytosolic ribosomes, represents a crucial process in maintaining mitochondrial function and activity. In order to test a direct effect of Aβ peptides on mitochondrial protein import, we utilized an *in vitro* assay system that measures the uptake of radiolabeled mitochondrial precursor proteins into intact mitochondria isolated from human cell cultures. This assay allows to directly follow the association, the uptake and the processing of mitochondrial precursor proteins (Ryan *et al.*, 2001; Chacinska *et al.*, 2009).

As precursor proteins, we used the following [^35^S]-labeled polypeptides: mitochondrial malate dehydrogenase (MDH2), an enzyme of the citric acid cycle; ornithine carbamoyltransferase (OTC) involved in the urea cycle; Su9(86)-DHFR and Su9(70)-DHFR, both artificial, mitochondrially targeted fusion proteins, comprising the presequence of the subunit 9 (Su9) of the F_1_F_0_-ATP synthase (86 and 70 AA respectively) from *Neurospora crassa* fused to the complete mouse dihydrofolate reductase (DHFR). All these precursor proteins contain an N-terminal presequence that is cleaved by the mitochondrial processing peptidase (MPP) after the polypeptide reaches the matrix compartment. Their mitochondrial import depends on the membrane translocase complexes TOM (<u>T</u>ranslocase of the <u>O</u>uter <u>M</u>itochondrial membrane) and TIM23 (<u>T</u>ranslocase of the <u>I</u>nner <u>M</u>itochondrial membrane with the core component Tim23) and a functional inner membrane potential (Δψ_mt_) (Chacinska *et al.*, 2009). In addition, we tested a precursor protein of the metabolite carrier family, the adenine nucleotide translocator 3 (ANT3). This protein is constituted by highly hydrophobic transmembrane subunits and lacks an N-terminal presequence. ANT3 is inserted into the inner mitochondrial membrane (IMM) and its import uses a distinct pathway that depends on the TOM and TIM22 complexes (Truscott *et al.*, 2002).

To assess AD-related pathological effects in the import assay, we used the most relevant Aβ peptides found in AD cases, constituted by 40 (Aβ40) and 42 (Aβ42) amino acids. The Aβ peptides and the radiolabeled precursor protein were incubated together with energized human mitochondria isolated from cultured HeLa cells. After the import incubation, samples were treated with proteases to digest residual non-imported polypeptides represented by the precursor form (*p*), and leaving the completely imported and processed mature form *(m*). Import reactions were analyzed by tricine-SDS-PAGE and Western blot followed by autoradiography to detect the ^35^S-labeled imported polypeptides, while the presence of Aβ peptides was detected by immunodecoration with a specific antibody against Aβ. As ANT3 does not contain a N-cleavable presequence and is not processed in the matrix, complete import was analyzed by blue-native gel electrophoresis (BN-PAGE) indicating the Δψ_mt_dependent formation of a dimeric complex after insertion into the inner membrane.

We found that Aβ peptides strongly interfered with the mitochondrial import of all precursor proteins analyzed (Figure *1*). The two Aβ peptides showed a different degree of inhibitory effect. Using the same concentration, Aβ40 partially inhibited the import reaction (Figure *1A*), while Aβ42 showed a complete inhibition (Figure *1B*) as indicated by the absence of the mature *(m*) form of a fully imported and processed precursor protein. ANT3 import was analyzed by BN-PAGE to visualize the formation of the complex around 150 kDa (Figure *1C*, lane *1*). Also in this case, Aβ peptides were able to inhibit the import reaction. Again Aβ42 was more effective in inhibiting the import reaction compared to Aβ40. The inhibitory effect, in particular of Aβ42, resulted in a full elimination of the generation of mature forms (m) as well as a complete protease sensitivity of the precursor protein (p) in the import reaction. Taken together, these two criteria indicate a full block of the mitochondrial translocation process and a general phenomenon affecting different import pathways.

**Figure 1.**
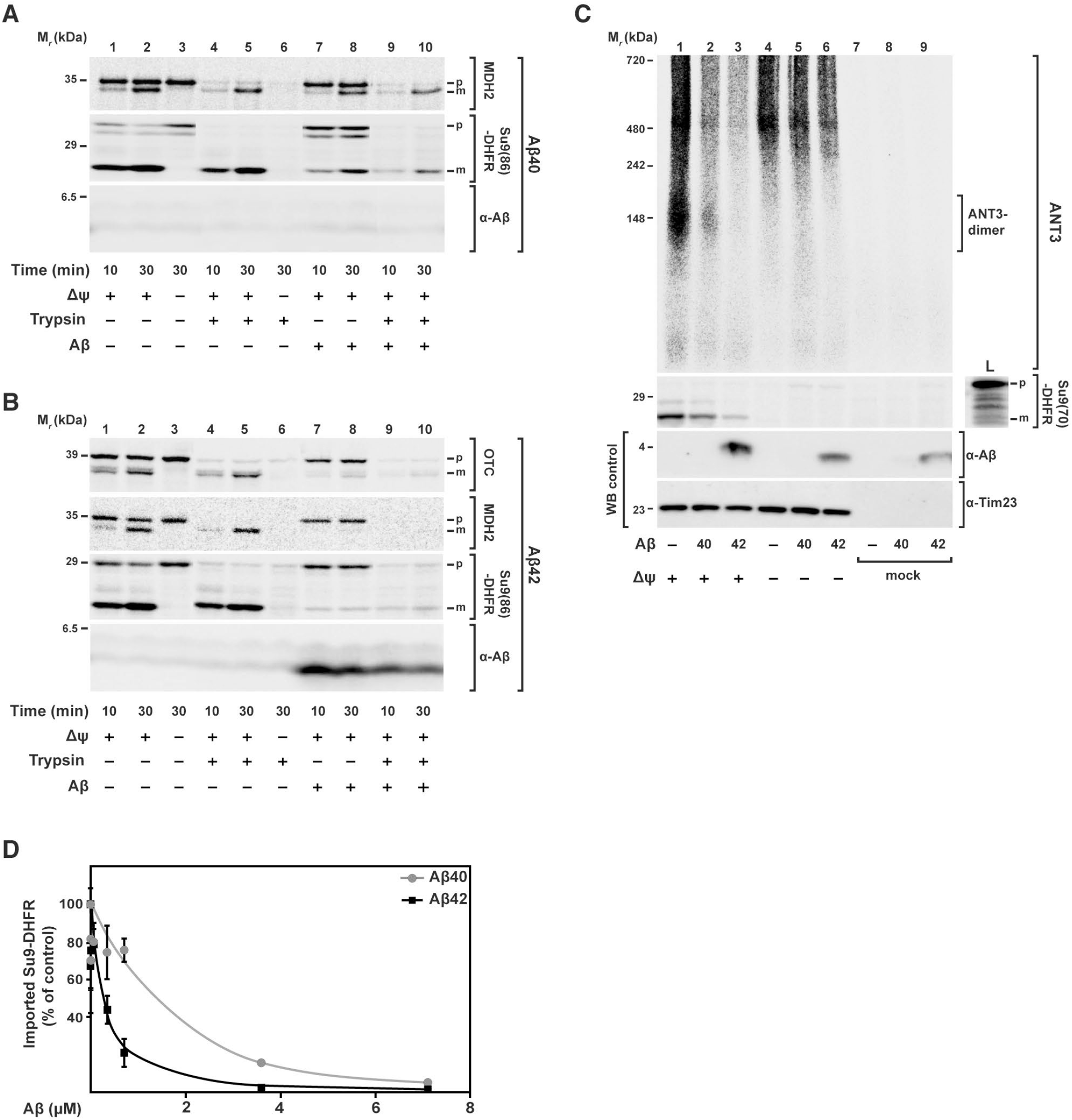
Effect of Aβ peptides on mitochondrial import of nuclear-encoded precursor proteins. [^35^S]-labeled radioactive precursor proteins were incubated with energized and isolated mitochondria from HeLa cell cultures in presence of same amounts (3.5 μM) of Aβ40 and Aβ42 peptides. (**A**, **B**) Import of the precursor proteins mitochondrial malate dehydrogenase (MDH2), the artificial reporter construct Su9(86)-DHFR, and ornithine carbamoyltransferase (OTC) for the indicated incubation times. After the import reaction, half of the samples (*lanes 4-6* and *9,10*) were treated with trypsin (100 μg/ml) to remove non-imported preproteins. Imported proteins were analyzed by tricine-SDS-PAGE followed by Western blot, digital autoradiography and immunodecoration against Aβ peptides. (**C**) Import of the adenine nucleotide translocator 3 (ANT3) in comparison with Su9(70)-DHFR. After import, all samples were treated with proteinase K (PK; 50 μg/ml) and analyzed either by BN-(ANT3) or SDS-PAGE (Su9(70)-DHFR), Western blot and digital autoradiography. As control, immunodecoration against Tim23 was carried out. (D) Quantification of import inhibitory effect of Aβ peptides. Import experiments with the precursor protein [^35^S]Su9(86)DHFR and different amounts of Aβ peptides (0.007 up to 7.0 μM) were performed as described above. The signals of processed and protease-resistant preprotein bands (*m-form*) were quantified using Image J. The amount of imported protein in the absence of Aβ peptide was set to 100%. Mean values and standard deviation (*S.D.*) were determined for n = 3 independent experiments. *p*, precursor protein; *m*, mature processed form; *L*, loading control; *WB.* Western blot.

In order to investigate the concentration-dependence of the inhibitory effect of Aβ peptides on mitochondrial import, we performed a titration of Aβ peptide amounts during the [^35^S]Su9(86)-DHFR import assay (Figure *1D* and Supplementary Figure *S1A* and *B*). After import, samples were digested by trypsin and analyzed by tricine-SDS-PAGE, autoradiography and Western blot. We quantified the protease-resistant mature form (*m*) of the imported [^35^S]Su9(86)-DHFR. We found that the inhibitory effect of Aβ42 was about ten fold stronger than Aβ40 (Figure *1D*). Inhibition of import by Aβ42 started at a concentration of about 0.1 μM, while for Aβ40 a concentration of more than 1 μM was required. It should be noted that only at the highest concentration, the Aβ40 band was detectable also in the mitochondrial fraction (Supplementary Figure *S1B*).

### Aβ peptides do not interfere with general mitochondrial functions

Since it was previously reported that *in vitro* Aβ peptides exert direct damage on mitochondria (Lustbader *et al.*, 2004; Hansson Petersen *et al.*, 2008; Mossmann *et al.*, 2014), we assayed the state of specific import-related mitochondrial functions in our experimental setup. An electric potential across the mitochondrial inner membrane (Δψ_mt_) is indispensable for the import of precursor proteins into the matrix as well as the insertion into the inner membrane (Ryan *et al.*, 2001). We measured the Δψ_mt_ in our model by the potential-dependent accumulation of the fluorescent dye tetramethylrhodamineethyl ester (TMRE) after incubation of isolated and energized mitochondria with increasing amounts of Aβ peptides (Figure *2A*). Both Aβ40 and Aβ42 did not exhibit any effect on Δψ_*m*__t_, even at high concentrations. As negative control, we incubated the mitochondria with 0.5 μM of valinomycin that causes a complete dissipation of the membrane potential and a concomitant strong reduction of the fluorescence signal. Using BN-PAGE, we inspected the structure and the composition of translocase complexes responsible for the import reaction under native conditions. In the BN-PAGE, the translocase complexes of both the outer membrane (TOM) and the inner membrane (TIM23) migrate as distinct high-molecular weight bands. Incubations with both Aβ peptides did not have any visible effect on the running behavior of the translocase complexes, indicating no significant change in structure and composition (Figure *2B*). Furthermore, the absence of effects in the native PAGE indicated that there is no significant stable interaction between the mitochondrial import complexes and Aβ peptides themselves. It should be noted that the detection of Aβ peptides in the BN gels (Figure *2B*), revealed a signal for Aβ42 localized in the upper part of the stacking gel, consistent with a formation of high molecular weight aggregates. Additionally, we also checked the running behavior of the five respiratory chain complexes of the inner membrane in native PAGE and again found no significant differences caused be the presence of Aβ peptides (Supplementary Figure *S2*). These results demonstrated that Aβ peptides did not negatively affect mitochondrial activities that are directly relevant for the import reaction. In line with this, resistance of mitochondrial control proteins against proteinase K (PK) treatment after import also suggests that mitochondrial membranes remained largely intact after Aβ treatment.

**Figure 2.**
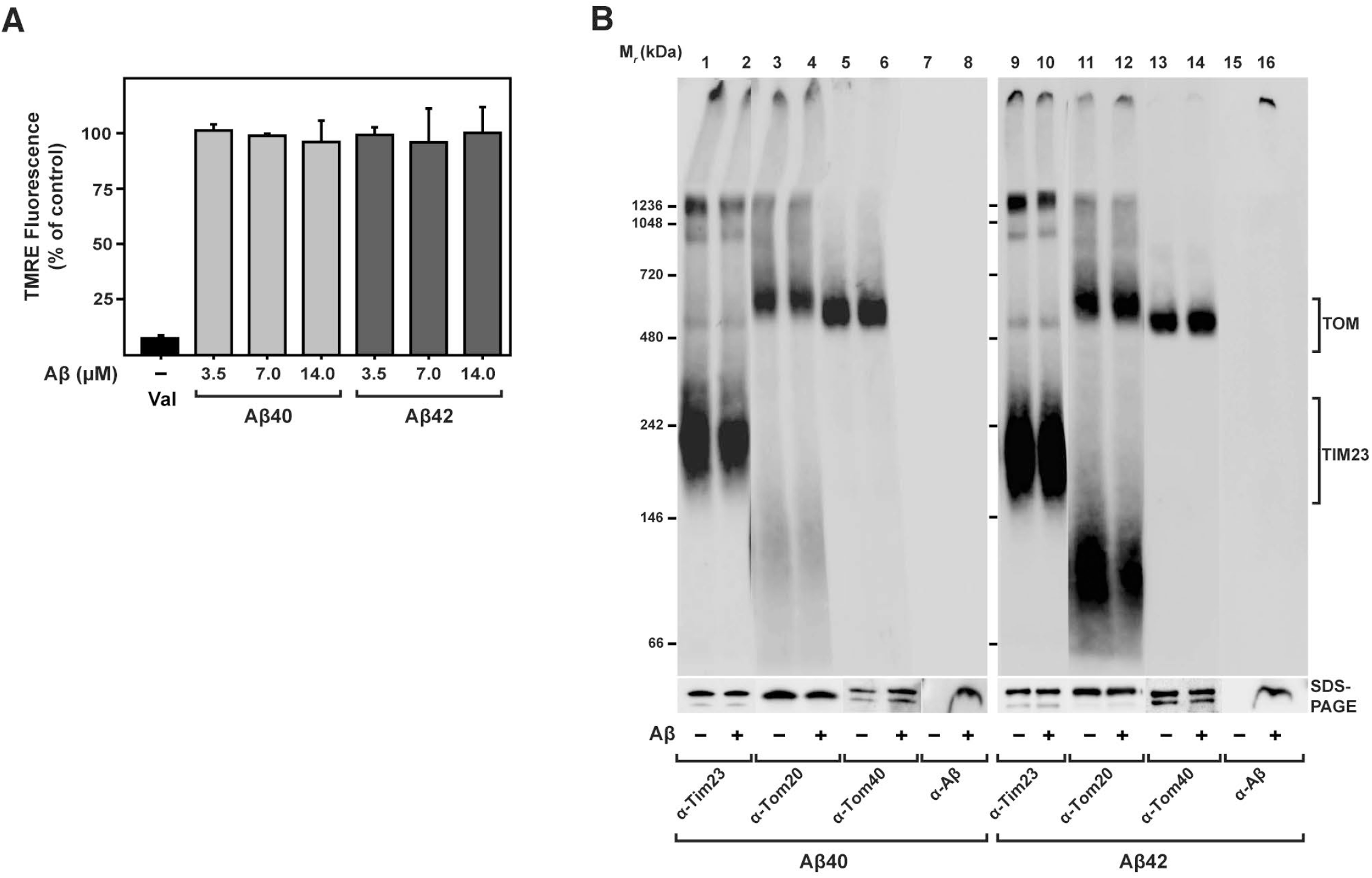
Effect of Aβ peptides on import-related mitochondrial functions. (**A**) Mitochondrial membrane potential (Δψ_mit_) was evaluated after treatment of energized mitochondria with increasing amount of Aβ peptides as indicated, followed by incubation with the potential-dependent fluorescent dye TMRE. After removal of excess TMRE, fluorescence was determined by a spectrofluorometer (Infinite M200 Pro, TECAN). Mean values and standard deviation were determined from three independent experiments. (**B**) After treatment of isolated and energized mitochondria with Aβ peptides (3.5 μM), structure and composition of import translocase complexes were analyzed by BN-PAGE, SDS-PAGE, and western blotting techniques. Before loading, mitochondria were solubilized in a buffer containing 1% digitonin. Immunodecorations against components of the translocase complexes TOM and TIM23, responsible for the import of presequence-containing preproteins through the mitochondrial membranes, Tom20, Tom40, Tim23 (*lanes 1-6* and *9-14*) and Aβ peptides (*lanes 7,8* and *15,16*) were performed.

### Aβ peptides affect the initial steps of the mitochondrial import reaction

Based on the observation of a significant inhibition of the overall import process, we set out to identify the particular step of the import reaction that was affected by Aβ peptides. Most cases of the precursor protein import can be generally distinguished into three steps: a) a binding to the receptors of the import machinery of the outer mitochondrial membrane (OMM); b) the Δψ_mit_-dependent transport through the membranes via the translocase complexes; c) the processing of the precursor to the mature form. To investigate the effect of Aβ peptides on the initial step of the import reaction, we dissipated the Δψ_mit_ as an import driving force, allowing only binding of precursor proteins to OMM import receptors and/or insertion into the TOM translocase channel. As the OMM binding reaction is very quick, we incubated the isolated mitochondria with the radioactive precursor protein for short times (range of seconds) in presence of Aβ peptides and tested for a co-fractionation of the precursor polypeptides with the mitochondria. Both Aβ peptides did not negatively affect the binding of the precursor protein [^35^S]Su9(86)-DHFR (Figure *3A*), indicating that the interaction with the mitochondrial surface receptors was not affected. On the other hand, in particular with Aβ42, we consistently observed elevated amounts of precursor protein associated with mitochondria that are proportional to the amount of peptide used (Supplementary Figure *S1C*). Since also non-specific radioactive protein bands, generated during *in vitro* translation in addition to the genuine precursor band, were found in association with the mitochondrial pellet after centrifugation, the increase in signal intensity of the precursor protein is probably due to an aggregation phenomenon (see below).

**Figure 3.**
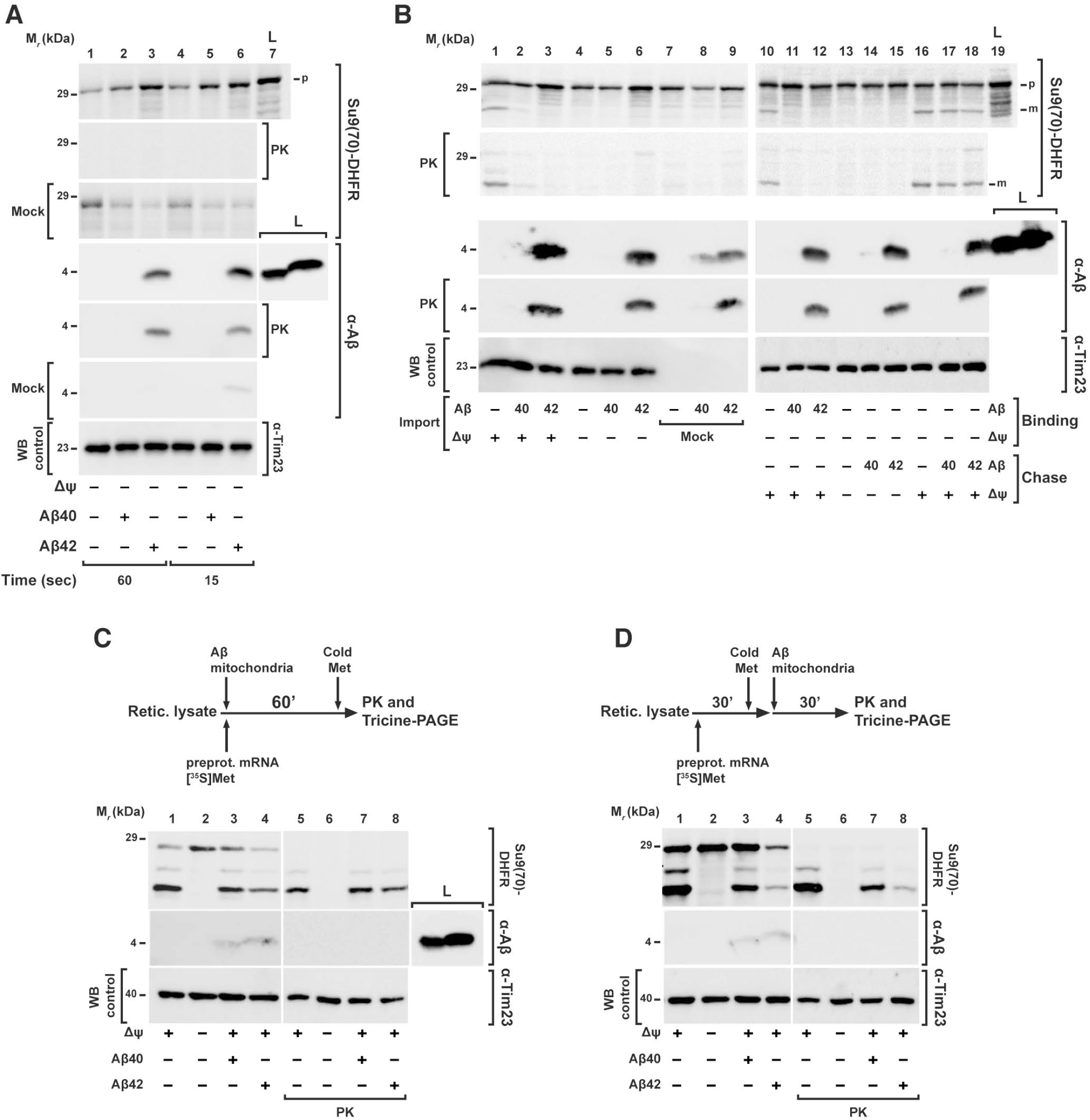
Mitochondrial import steps affected by Aβ peptides. (**A**) Binding of the precursor protein to the OMM import machinery receptors. After removing the Δψ_mit_, mitochondria were incubated for short time points (range of seconds) with Aβ peptides (3.5 μM) and precursor protein [^35^S]Su9(70)DHFR. Half of the samples were incubated with proteinase K (PK; 50 μg/ml) to digest not imported precursor protein. (**B**) Separation of preprotein binding (*Binding*) to OMM from inner membrane translocation and processing steps (*Chase*). For precursor binding and insertion into the OMM, Δψ_mit_ was dissipated by CCCP (1 μM) during incubation with [^35^S]Su9(70)DHFR in presence (lanes *11,12*) and absence of Aβ peptides (lanes *10* and *13-18*). To assay inner membrane translocation and processing (*Chase*), the Δψ_mit_ was restored by addition of albumin (BSA; 2 mg/ml; lanes *10-12* and *16-18*) in presence (lanes *17,18*) and absence of Aβ peptides. For comparison, a complete one-step import reaction was performed (lanes *1-9*). (**C, D**) Effect of Aβ peptides on co- and post-translational import. (**C**) As shown in the scheme of the experimental setup, a co-translational import was performed by incubating rabbit reticulocyte lysate, Su9(70)DHFR mRNA, [^35^S]Methionine, and isolated energized mitochondria in presence or absence Aβ peptides (3.5 μM) as indicated at 30° C for 60 min. (**D**) In the post-translational import, first the translation of [^35^S]Su9(70)DHFR was performed using rabbit reticulocyte lysate, Su9(70)DHFR mRNA, [^35^S]methionine for 30 min, and then isolated mitochondria were added in presence of Aβ peptides (3.5 μM) for additional 30 min to perform the mitochondrial import reaction. The translation was stopped by adding 8 mM cold methionine. All samples were analyzed by tricine-SDS-PAGE followed by Western blot, digital autoradiography and immunodecoration against Aβ peptides and Tim23. *p*, mitochondrial precursor protein; *m*, mitochondrial mature form; *Mock*, control experiment in the absence of mitochondria; *L*, total amount of Aβ peptides added*. WB*, Western blot.

Effects on the transport and processing reactions were tested by a two-step protocol that separated the binding of the precursor from the actual translocation process. The precursor protein [^35^S]Su9(70)-DHFR was first incubated with mitochondria where the Δψ_mit_ was dissipated by the addition of CCCP (1 μM). In this way, the precursor protein was able to bind to the TOM machinery without being imported. After removing excess unbound precursor proteins, Δψ_mit_ was restored by taking away the CCCP by binding it to excess amounts of albumin (BSA) and re-energizing the mitochondria, allowing the translocation and processing reaction to proceed. Interestingly, an inhibition of protein import was only observed when Aβ peptides were present already in the first step of the experiment, (Figure *3B*, lanes *11* and *12*). While adding the peptides directly in the second step, after the binding step has been completed, did not show any effect on the import reaction (Figure *3B*, lanes *17* and *18*). This directly demonstrated that Aβ peptides did not negatively affect the later phases of the import reaction, but rather interfered with the first steps of the import reaction that happen at the outer face of the OMM.

In the used *in vitro* system, the translation reaction to produce radiolabeled preproteins is separated from the actual import reaction, essentially resulting in a post-translational translocation process. However, in cells, the mitochondrial import most likely represents a mixture of post- and co-translational reactions, depending on the individual properties of preproteins or even their mRNAs (Fox, 2012). To clarify if Aβ peptides were able to inhibit mitochondrial import also during a co-translational mechanism, we decided to perform a import reaction in the reticulocyte lysate system used for producing the [^35^S]-labeled preprotein Su9(70)-DHFR. We incubated the reticulocyte labeling mix, containing ribosomes, energized mitochondria, preprotein mRNA, and [^35^S]-methionine, for 60 min in presence or absence of Aβ40 peptides (Figure *3C*). After analyzing the samples by tricine-SDS-PAGE, we did not observe an import inhibition in presence of Aβ40 but still a significant reduction of the formation of the mature form in presence of Aβ42, although not as pronounced as in the post-translational situation. As a post-translational control, we used the same experimental setup but first performed the translation reaction without mitochondria for 30 min, stopped the labeling by addition of non-labeled (“cold”) methionine and only then added the isolated energized mitochondria in presence or absence of Aβ peptides (Figure *3D*). In this case, we observed a partial inhibition of mitochondrial import in presence of Aβ40, while Aβ42 gave a strong inhibitory effect. The less severe inhibitory effect in case of a forced co-translational import situation support our conclusion that Aβ peptides do not directly affect mitochondria function but rather act at a more initial step of the import process. Off note, in a control experiment without mitochondria, we observed that Aβ peptides were not affecting ribosomal translation rates (Supplementary Figure *S1D*).

### Interaction of Aβ peptides with human mitochondria

As our experiments indicated the possibility of a direct association of Aβ peptides, in particular Aβ42, with mitochondria, despite any obvious deleterious effects on mitochondrial functions, we set out to analyze the inhibition properties on preprotein import in more detail. Since the degree of import inhibition seemed to correlate with the amount of Aβ peptides co-purified with mitochondria, we checked if Aβ peptides could also act in absence of precursor proteins. We pre-treated isolated mitochondria with Aβ peptides for 30 min followed by several washing steps to remove excess unbound material. Then, we performed a normal import reaction using the precursor protein [^35^S]Su9(86)-DHFR without peptide addition (Figure *4A*). Interestingly, a pre-treatment of mitochondria with Aβ40 did not show any significant co-purification of Aβ40 with the mitochondria and also did not affect a later import reaction (Figure *4A*, lanes *9, 12* and *15*). On the contrary, a pretreatment of mitochondria with Aβ42 showed a strong, although not complete, inhibitory effect on the subsequent import reaction. Furthermore, we were able to detect Aβ42 co-purifying with the mitochondria even after extensive washing, confirming an association with mitochondria (Figure *4A*, lanes *10, 13* and *16*).

**Figure 4.**
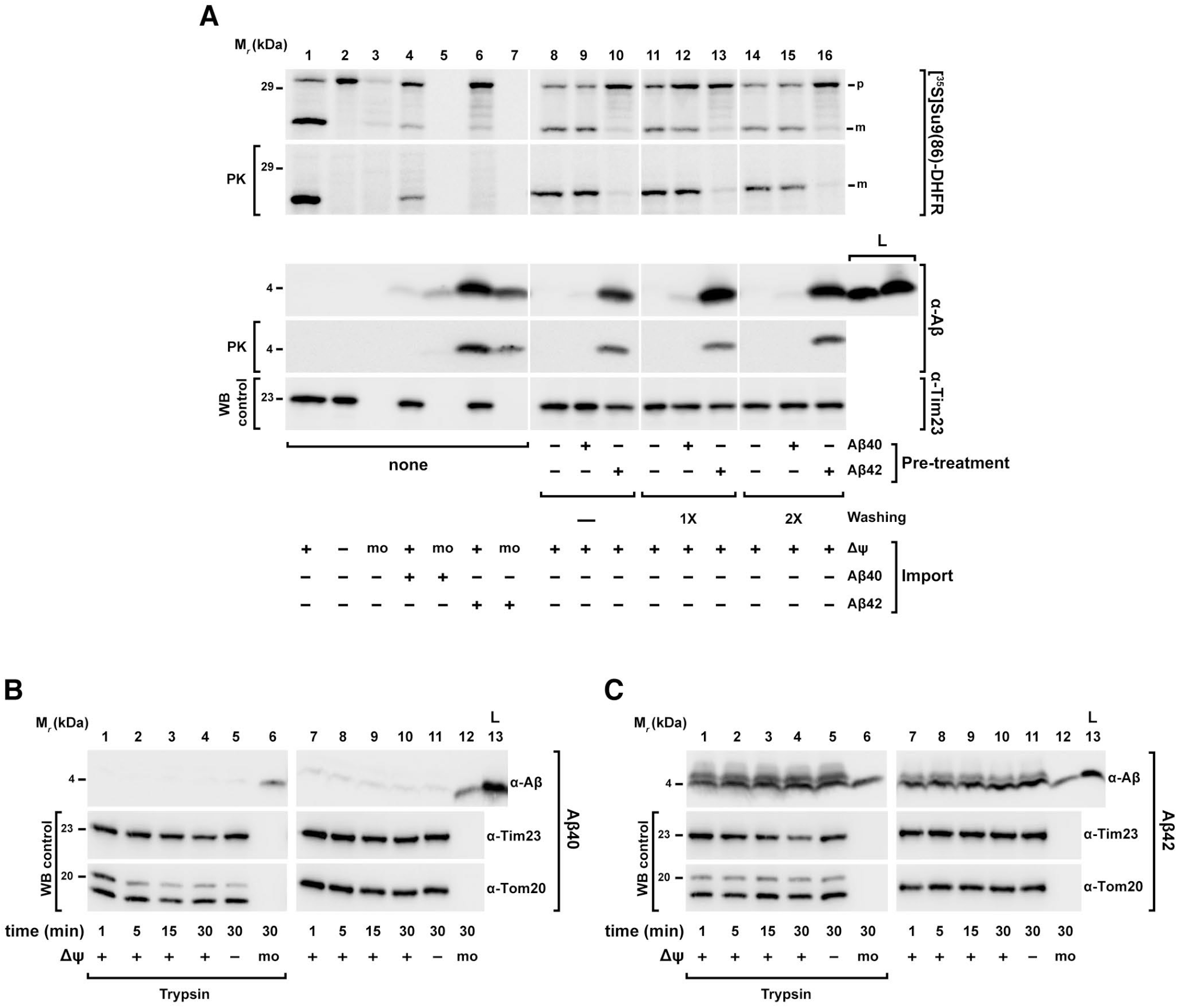
Analysis of Aβ peptides interaction with human mitochondria. (**A**) Pretreatment of mitochondria with Aβ peptides. Isolated mitochondria were incubated with Aβ peptides (3.5 μM) at 30° C for 30 minutes. After several washing steps, mitochondria were re-isolated and incubated in an energizing buffer with precursor protein [^35^S]Su9(86)DHFR for an import reaction in the absence of Aβ peptides (*lanes 8-16*). For comparison, the precursor protein [^35^S]Su9(86)DHFR was directly incubated with isolated and energized mitochondria and in mock samples (*mo*) in presence or absence of Aβ peptides (*lanes 1-7*). Half of the samples were treated with proteinase K (PK; 50 μg/ml) to digest not imported precursor protein. (**B, C**) Mitochondrial association of Aβ peptides. Isolated and energized mitochondria and mock (*mo*) samples (*lanes 6*, *12*) were incubated with the same amount of Aβ40 (**B**) and Aβ42 (**C**) peptides (3.5 μM) for different time points. Δψ_mit_ was dissipated where indicated (*lanes 5 and 11*). Half of the samples were then treated with trypsin (100 μg/ml; *lanes 1-6*). All samples were processed by tricine-SDS-PAGE followed by Western blot. As control, immunodecorations against mitochondrial Tom20, Tim23 and Tom40 proteins were performed. *WB*, Western blot; *p*, precursor protein; *m*, mature form.

We therefore investigated the biochemical properties of an association of Aβ peptides with isolated mitochondria more thoroughly. First, we performed a standard mitochondrial import experiment using Aβ peptides to clarify if they were taken up via the canonical import pathway. The import reaction was analyzed by tricine-SDS-PAGE followed by Western blot using antiserum against Aβ peptides. As shown in Figure *4B*, the smaller peptide Aβ40 again did not show a significant co-purification with mitochondria even at longer incubation times. In contrast, with Aβ42, as seen before, a band of 4 kDa was visible in the samples containing mitochondria already at very short time points (Figure *4C*). The band intensity did only slightly increase with longer incubation times. Due to the small size and the specific properties of the Aβ peptides, any processing event during a potential import reaction was not expected. However, for Aβ42 an additional band with a slightly higher molecular weight appeared in the presence of mitochondria, which is likely due to a different running behavior of the small peptide in presence of high amounts of mitochondrial proteins or lipids. However, three observations argue strongly against a specific uptake of Aβ peptides via the mitochondrial import machinery: a) no time-dependence of the mitochondria-associated signals (e. g. Figure *4C*, lanes *7-10*), b) the intensity of the co-purifying Aβ signal was not influenced by Δψ_*mit*_ (Figure *4C*, lane *11*), and c) both Aβ peptides showed a comparable signal also in the mock sample containing no mitochondria at all (Figure *4B* and *C*, lanes *6* and *12*). Interestingly, both the co-purifying materials as well as the peptides in the mock samples were largely resistant to protease digestion (Figure *4B* and *C*, lane *6*).

As protection against proteases is a major hallmark of a successful mitochondrial import reaction (Ryan *et al.*, 2001), we characterized the protease digestion behavior of Aβ peptides in more detail (Figure *5A*). We incubated the Aβ peptides with isolated and energized mitochondria followed by solubilization with 0.5% Triton X-100 (Figure *5A*, lanes *5-8*) or ultra-sonicatio (Figure *5A*, lanes *9-12*). Under these conditions, the mitochondrial membranes are disrupted and would not be able to offer protection against external proteases. A titration with rising amounts of trypsin was performed and then all the samples underwent trichloroacetic acid (TCA) precipitation, tricine-SDS-PAGE and detection of present Aβ peptides by Western blot. As showed in control panels, both detergent- and sonication-lysis of mitochondria were successful as endogenous control proteins were efficiently degraded even at the lowest concentration of trypsin (5 μg/ml). In the mock samples, without mitochondria and used as control, we again found a significant protease resistance of both Aβ peptides (Figure *5A*, lanes *1-4*). The protease resistance of both Aβ peptides was decreased in presence of detergent or after ultrasound treatment (Figure *5A*, lanes *6-8* and *10-12*). Aβ42 was found slightly more resistant than Aβ40 after detergent lysis, but remained resistant to trypsin after ultrasound treatment. In presence of mitochondria, the behavior of the two peptides was different. As Aβ40 did not co-purify or pellet with mitochondria, the analysis of Aβ40 susceptibility to protease digestion was not possible. In contrast, Aβ42 showed some co-purification with the mitochondria and also a complete protease resistance that was neither affected by the presence of detergent nor by sonication. This intrinsic protease resistance and the detection of Aβ peptides in samples without mitochondria (mock) or after destruction of mitochondrial membranes suggest that in our experimental setup Aβ peptides are more prone to form sedimentable aggregated material than to associate with the OMM. We also investigated if the intrinsic resistance of Aβ peptides to protease digestion was specific only for trypsin or could be extended to other proteases. We incubated equal amounts of Aβ peptides alone or with isolated mitochondria for 30 min at 30˚ C and subsequently treated all samples with the proteases trypsin or proteinase K (PK) for different times, followed by tricine-SDS-PAGE and Western blot (Figure *5B*). As observed already before, Aβ40 was not co-purifying with the mitochondria. However, in mock samples, Aβ40 was partially resistant to both proteases even after longer treatments. Aβ42 partially associated with mitochondria, but was not completely digested by both proteases (Figure *5B*, lanes *5-12*), in mitochondria as well as mock samples, indicating that both Aβ peptides retain an intrinsic protease resistance that is independent of a mitochondrial association.

**Figure 5.**
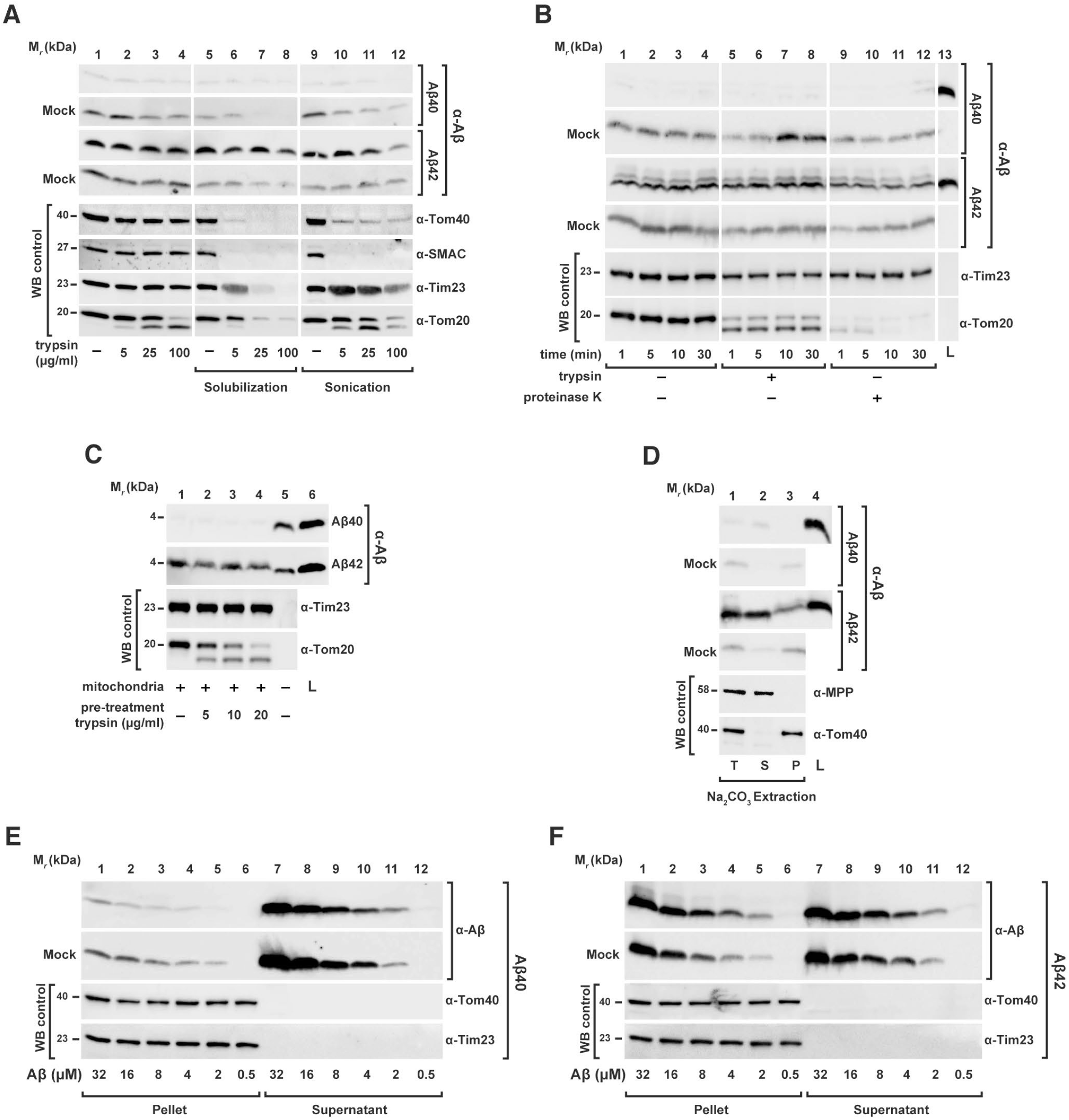
Membrane interaction behavior of Aβ peptides. (**A**) Intrinsic protease resistance. Aβ peptides (3.5 μM) were incubated with or without (*Mock*) intact and energized mitochondria followed by digestion with increasing amounts of trypsin (*lanes 1-4*). As controls, mitochondria were lysed by solubilization with 0.5% Triton X-100 (*lanes 5-8*) or by sonication (*lanes 9-12*) before the addition of the protease. All the samples underwent TCA precipitation. (**B**) Protease sensitivity. Aβ peptides (3.5 μM) were incubated with or without (*Mock*) intact and energized mitochondria followed by digestion with 100 μg/ml trypsin (*lanes 5-8*) and 100 μg/ml proteinase K (PK) for different time points as indicated in the figure. (**C**) Dependence of the interaction between Aβ peptides and isolated mitochondria on peripheral OMM receptors. Isolated mitochondria were pre-treated with the indicated trypsin concentrations to digest exposed OMM proteins. After trypsin inactivation, isolated mitochondria were re-isolated and incubated in an energized buffer with Aβ peptides (3.5 μM). (**D**) Alkaline extraction of Aβ peptides from mitochondria and mock samples. Aβ peptides (3.5 μM) were incubated in presence or absence (*Mock*) of isolated and energized mitochondria. After reisolation, mitochondria and mock samples were subjected to alkaline extraction as described under “Material and Methods” section. (**E, F**) Titration of Aβ peptide amounts. Increasing concentrations of Aβ40 (**E**) and Aβ42 (**F**) peptides were incubated for 30 min in presence or absence (*Mock*) of energized mitochondria and separated in insoluble (*Pellet*) and soluble (*Supernatant*) fractions. All samples were analyzed by tricine-SDS-PAGE and Western blot. As control, immunodecoration against the endogenous mitochondrial proteins such as SMAC (IMS), MPP (matrix) and Tom40 (OMM) was carried out. *T*, total; *P*, pellet; *S*, supernatant; *L*, loading control; *WB*, Western blot.

The import of nuclear-encoded precursor proteins initially requires a specific interaction with receptor proteins at the surface of the OMM (Endo and Kohda, 2002). To analyze if the interaction of Aβ peptides with mitochondria depends on the involvement of the OMM receptors, we pre-treated isolated intact mitochondria with trypsin to digest any protein domains exposed on the cytosolic face of the outer membrane. Then, we incubated the mitochondria with Aβ peptides (Figure *5C*). Samples were analyzed by tricine-SDS-PAGE followed by Western blot. As control, Tom20 was degraded at the lowest trypsin concentration (5 μg/ml), while the inner membrane protein Tim23 was stable during both protease treatments indicating the intactness of mitochondria. The co-purified amount of Aβ42 with mitochondria did not show any difference between trypsin pre-treated mitochondria versus untreated control samples, indicating that any potential interaction of Aβ42 with mitochondria is not based on a specific binding to the import-related receptor proteins of the TOM complex.

The previous experiments suggest that the association of Aβ peptides with mitochondria rather represents a non-specific interaction with the OMM. We performed an alkaline extraction to assess the membrane interaction properties after incubating Aβ peptides with mitochondria, (Figure *5D*). During alkaline extraction, polypeptides that stably associate with membranes remain in the pellet fraction (*P*), while peripheral membrane proteins are found in the supernatant (*S*). As shown before, Aβ40 did not show a significant signal in presence of mitochondria. However, the mock samples showed that minor amounts of Aβ40 accumulated in the pellet fraction consistent with a generation of small amounts of protein aggregates. The Aβ42 peptides showed a similar behavior in the mock samples. However, in the presence of mitochondria, a significant amount of co-purified material was found in the supernatant fraction excluding integration into the OMM, suggesting at most a peripheral association. The mitochondrial control proteins MPP (soluble) and Tom40 (membrane-integrated) behaved as expected. A non-specific interaction with the OMM, in particular for Aβ42, was also supported by a saturation titration experiment (Figure *5E* and *F*). Here, we incubated increasing amounts of Aβ peptides with a constant amount of mitochondria and separated soluble and insoluble material by intermediate-speed centrifugation. Increasing the peptide concentration, most of the Aβ40 peptide remained in the supernatant and only a minor amount appeared in the pellet fraction (Figure *5E*) without being influenced by the presence of mitochondria. On the other hand, significant amounts of Aβ42 peptides accumulated in the pellet fraction, both in presence or absence of mitochondria (Figure *5F*). In both cases, the amount of Aβ42 peptides recovered in the pellet fractions did not seem to be saturable, indicating again a non-specific mitochondrial association as well as a pronounced tendency to form sedimentable aggregate material.

From the results above, it was not possible to clearly distinguish between Aβ peptides associated to the OMM and Aβ peptides prone to aggregation that are able to sediment with mitochondria by conventional differential centrifugation methods used in a standard import assay. Thus, we decided to analyze the behavior of Aβ peptides during the mitochondrial import using a specific rate-zonal centrifugation method. After performing an import reaction with the precursor protein [^35^S]Su9(70)-DHFR in presence or absence of Aβ peptides, samples were separated by centrifugation through a sucrose gradient (20-50%). Fractions from top to bottom were collected and analyzed by Western blot or autoradiography for the presence of the imported precursor protein or Aβ peptides (Figure *6*). As controls, we carried out the same experiment in the absence of mitochondria (mock) or in the absence of Aβ peptides (Figure *6B*). From the sedimentation behavior of mitochondrial marker MPP and Tim23, isolated mitochondria were concentrated mostly around the middle of the gradient (Figure *6E*, fractions *12-14*). Most of Aβ40 accumulated as monomer or as small, low density and SDS-soluble aggregates at the top of the gradient and no co-sedimentation with the mitochondria was observed (Figure *6A*, upper panels). This observation is consistent with the behavior in our differential centrifugation experiments (see above). However, Aβ42 behaved significantly different (Figure *6A*, middle panels). In presence of isolated mitochondria, a small percentage of Aβ42 (20% of the total Aβ42 added to the experiment; see Figure *6E*) was found in the gradient fractions together with the mitochondrial markers, suggesting a direct interaction with mitochondria. In the mock samples, most of Aβ42 accumulated on the top of the gradient like Aβ40. In the control samples containing only the precursor protein, [^35^S]Su9(70)-DHFR showed a localization of the mature form (*m*) in the same fractions as the bulk mitochondria (Figure *6B*). As expected, in presence of Aβ40 the amount of mature form was partially reduced (Figure *6C*), while Aβ42 treatment resulted in a complete disappearance of the mature form, demonstrating again a complete inhibition of mitochondrial import (Figure *6D*). Interestingly, the presence of precursor proteins changed the behavior of Aβ42 as the amount of mitochondria-associated material decreased while the amount in the bottom fractions, representing aggregates increased (Figure *6D* and *E*). Additionally, in the presence of Aβ42 a considerable amount of the precursor protein itself was found in the aggregate fraction at the bottom of the gradient, indicating the formation of co-aggregates between Aβ peptides and mitochondrial precursor proteins (Figure *6D*, lane 23 and Figure *6E*).

**Figure 6.**
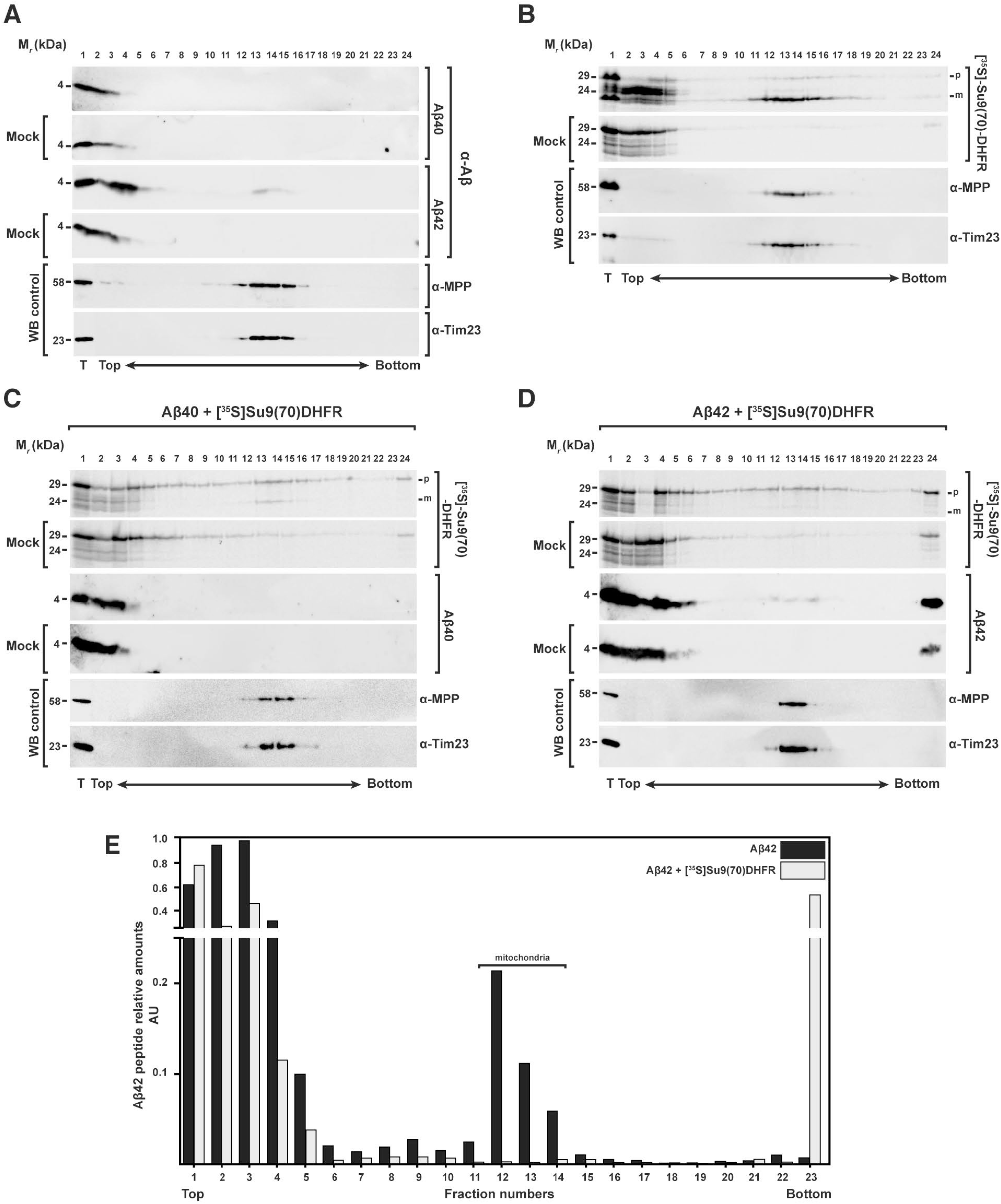
Analysis of the interaction between Aβ peptides and mitochondrial precursor proteins with mitochondria through density gradient centrifugation. (**A**) Sucrose gradient centrifugation of 3.5 μM Aβ40 (upper panels) and Aβ42 (lower panels) incubated with and without (Mock) isolated and energized mitochondria. (**B**) As control, a sucrose gradient of precursor protein [^35^S]Su9(70)DHFR incubated with or without (*Mock*) isolated and energized mitochondria in the absence of Aβ peptides was performed. (**C, D**) Sucrose gradients with or without (*Mock*) mitochondria incubated with precursor protein [^35^S]Su9(70)DHFR in the presence of Aβ40 (**C**) or Aβ42 (**D**). Density gradient fractionations were performed as reported in “Materials and Methods” section. Samples were analyzed by tricine-SDS-PAGE and Western blot. As control, immunodecorations against MPP and Tim23 were used. (**E**) Quantification of the Aβ42 band intensities incubated with mitochondria in absence (**A**) or presence (**D**) of precursor protein [^35^S]Su9(70)DHFR. Each value is the ratio between the intensity of the Aβ42 band in each fraction and the total sample (*T*). *WB*, Western blot; *p*, precursor form; *m*, mature form of the preprotein.

Taken together these data do not support previous conclusions about an uptake of Aβ peptides into the organelle as the typical criteria for mitochondrial import were not fulfilled: specificity for mitochondria, dependence on surface import receptors and Δψ_mit_, and acquisition of protease resistance. However, we observed some degree of non-specific association with the mitochondrial surface in case of Aβ42. This peptide also exhibited a strong tendency to form aggregates, independently of the presence of mitochondria. Interestingly, in presence of mitochondrial precursor proteins, the association of Aβ42 with the mitochondria was reduced while at the same time an increased formation of sedimentable preprotein-Aβ42 conglomerates was observed.

### Preprotein import competence is reduced by the formation of Aβ-preprotein coaggregates

As aggregate formation is an intrinsic pathological property of Aβ peptides (Thal *et al.*, 2015), we reasoned that a reduction of preprotein solubility by aggregation in presence of Aβ peptides might contribute to the inhibitory effect on the import reaction. We therefore analyzed a co-aggregation by three types of assays: a) high-speed centrifugation followed by tricine-SDS-PAGE, b) filter retardation assay, and c) BN-PAGE. These techniques provide direct information about the aggregation behavior of precursor polypeptides in the presence of the Aβ peptides and partially characterize the nature of the aggregates. After incubation of radiolabeled precursor proteins with Aβ peptides, samples were centrifuged at high speed (45.000 rpm; 124.500 xg) to separate the insoluble high-molecular weight aggregates from the soluble proteins. The resulting pellets and supernatants were analyzed by Western blot and immunodecoration against Aβ peptides, as well as autoradiography to detect the precursor polypeptides (Figure *7A*). The precursor protein alone partially fractionated to the pellet suggesting an intrinsic aggregation propensity (Figure *7A*, lanes *7* and *17*). However, in presence of rising concentrations of Aβ42, the amounts of [^35^S]Su9(86)-DHFR found in the pellet was significantly increased (Figure *7A*, lanes *18-20*). In contrast, Aβ40 had less severe effects on the distribution of precursor polypeptides in the centrifugation assay (Figure *7A*, lanes *8-10*) where most precursor protein remained soluble in the supernatant (Figure *7A*, lanes *3-5*). Aβ42 itself was mostly found in the pellet fraction, suggesting a strong propensity to form insoluble aggregates (Figure *7A*, lanes *16, 18-20*). In the pellet fraction, but not in the supernatant, an additional band was detected for Aβ42 at the top part of the tricine gel corresponding to the loading pockets. This suggested that Aβ42 formed high-molecular weight aggregates that were insensitive to SDS solubilization. For Aβ40, part of the peptides sedimented as insoluble aggregates (Figure *7A*, lanes *6,8-10*) and part remained soluble in the supernatant (Figure *7A*, lanes *1,3-5*). In the supernatant fraction, Aβ40 showed two bands around 20 kDa and 35 kDa in addition to to the predominant band at 4 kDa (Figure *7A*, lanes *3* and *4*). These bands were present only when Aβ40 was incubated with the precursor proteins, but not with the peptides alone. Similar bands were also detected with Aβ42, but in much lower amounts (Figure *8A*, lanes *12* and *13*).

**Figure 7.**
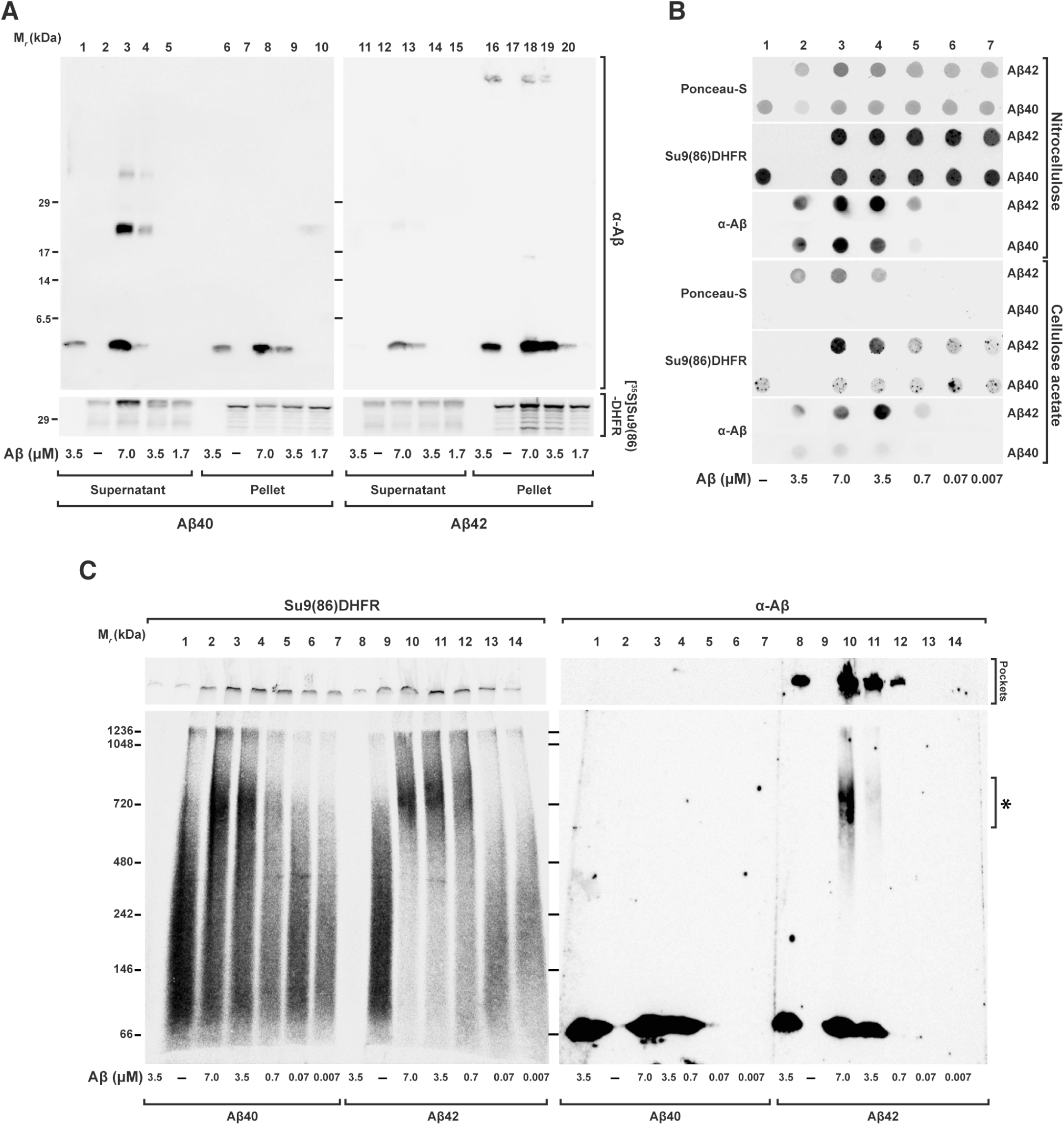
Co-aggregation between Aβ peptides and mitochondrial precursor protein. Precursor protein [^35^S]Su9(86)-DHFR was incubated for 30 min at 30 °C in import buffer in presence or absence of the indicated amounts of Aβ peptides. After incubation, samples were analyzed by the following techniques: (**A**) Tricine-SDS-PAGE. Soluble fractions (*Supernatant*) were separated from the insoluble (*Pellet*) by centrifugation for 40 min at 123000 xg and 4 °C. Samples were analyzed by tricine-SDS-PAGE. (**B**) Filter retardation assay. Samples were filtered directly through cellulose acetate and nitrocellulose membranes using a dot blot filtration unit as described in “Material and Methods” section. Proteins bound to both membranes were stained with Ponceau S. Bound Aβ peptides were detected by immunodecoration and the precursor protein by digital autoradiography. (**C**) BN-PAGE. Samples were loaded on native PAGE as described in “Materials and Methods” and analyzed by Western blot. The precursor protein signal was detected by digital autoradiography and the Aβ peptides by immunodecoration.

In the filter retardation assay, different amounts of Aβ peptides were incubated with the [^35^S]Su9(86)-DHFR (Figure *7B*) or [^35^S]OTC (Supplementary Figure *S3*) and subsequently filtered through nitrocellulose or cellulose acetate membranes. With the cellulose acetate membrane, which does not have an intrinsic protein binding affinity, inclusions or aggregates bigger than 0.2 μm are trapped under these conditions, while the smaller complexes pass through and are washed away (Heiser *et al.*, 2000). As most of the added protein should be retained on a nitrocellulose membrane, this type of membrane was used as loading control. Precursor proteins were detected by autoradiography and the presence of Aβ peptides by immunodecoration. The total amount of retained polypeptides was also evaluated by Ponceau red staining of the membranes. As expected from their intrinsic aggregation propensities, Aβ42, but not Aβ40, showed a signal on cellulose acetate membranes, when similar concentrations were loaded (Figure *7B*). While the precursor protein [^35^S]Su9(86)-DHFR alone showed a light signal on cellulose acetate membrane, a strong signal was detected when it was incubated together with Aβ42 (Figure *7B*). The formation of the precursor protein aggregates increased with the amount of Aβ42 peptides added. [^35^S]OTC showed a similar behavior (Supplementary Figure *S3*).

We also applied the samples on BN-PAGE to characterize the complex formation between Aβ peptides and precursor proteins under native condition. After incubation of the [^35^S]Su9(86)-DHFR with different concentrations of Aβ peptides, the complete samples were separated by BN-PAGE gradient gel (5-16.5%) and then analyzed by Western blot and autoradiography. The precursor protein [^35^S]Su9(86)-DHFR alone distributed over a large size range without forming a defined band, a typical behavior for a soluble protein in native PAGE (Figure *7C*, lanes *2* and *9*). In presence of Aβ40, some of the precursor proteins shifted to a higher molecular weight zone of the gel in a concentration-dependent manner (Figure *7C*, lanes *3-7*). In presence of Aβ42, the signals of the precursor protein almost exclusively shifted to an area around 720 kDa (Figure *7C*, lanes *10-13*). Interestingly, the immunodecoration with anti-Aβ serum showed that some Aβ42 material accumulated at the same molecular weight range (Figure *7C*, lanes *10* and *11*). In addition, Aβ42 also exhibited a signal at the highest part of the membrane related to the loading pockets in the gel, representing large insoluble aggregate material (Figure *7C* lanes *8, 10-12*). The fact that in native conditions the precursor protein band together with Aβ42 band shifted to the same area strongly suggests a direct interaction between the precursor protein and Aβ42. The large size of the complex, comprising multiple copies of both molecules was consistent with the formation of Aβ42-preprotein co-aggregates.

Taken together, the data obtained from three different technical approaches clearly confirmed a co-aggregation phenomenon between the precursor proteins and Aβ peptides that reduced the precursor proteins solubility. As solubility of the precursor proteins is a requirement for an efficient mitochondrial import, a formation of co-aggregates between the precursor proteins and Aβ peptides interferes with the insertion of the precursor protein inside the TOM channel. This represents the initial step of an import reaction that was found defective in our experiments in presence of Aβ peptides. Notably, the two Aβ peptides analyzed showed different effects on co-aggregate formation, correlating well with the observed preprotein inhibition efficiency, their aggregation propensity and also the pathological impact in AD patients.

## Discussion

Pathological properties of intracellular Aβ peptides, in particular in correlation with mitochondrial dysfunction have been previously observed on many occasions in the context of AD. Aβ peptides a) localize to mitochondria from postmortem AD brains and from several experimental models of the disease (Pagani and Eckert, 2011), b) physically interact with some mitochondrial components (Lustbader *et al.*, 2004), and c) exert harmful effects on mitochondrial function (Kaminsky *et al.*, 2015). As an *in situ* production of Aβ peptides in mitochondria themselves seems unlikely (Sannerud and Annaert, 2009), our study addressed the possible mechanisms of Aβ peptide interaction with mitochondria as well as the correlation between a mitochondrial localization of Aβ peptides and the mitochondrial dysfunctions observed in AD.

Under *in vitro* conditions, we observed a clear-cut and strong inhibitory effect of Aβ peptides on mitochondrial import. The inhibitory effect of the Aβ42 was significantly stronger than the related Aβ40, correlating well with the stronger pathogenic effect of Aβ42 in human AD patients (Eckman and Eckman, 2007). Notably, the Aβ42 concentration that resulted in a significant inhibition of mitochondrial import was comparable to the concentration of the peptide that has previously found in AD brains, 2 μM for Aβ42 (Roher *et al.*, 2009). Our experiments also shed a light on the biochemical details of the inhibitory mechanism, in particular which stage of the import process was affected. The inhibitory effect occurred immediately and did not require a prolonged preincubation period. Although previous publications reported that a treatment of mitochondria with Aβ peptides resulted in a reduction of the Δψ_mit_ (Kaminsky *et al.*, 2015), in our model system we did not observe any changes in Δψ_mit_ in the time-frame of the import experiments, excluding an Aβ-related reduction of the membrane potential as a cause for the import inhibition. Neither did we observe changes in the size and composition of the precursor protein translocase complexes (TOM and TIM) nor of the metabolic complexes of the respiratory chain. The possibility of a direct physical damage on mitochondrial membranes, the oxidative phosphorylation system or the preprotein import machinery by Aβ peptides is therefore very unlikely.

Most of the mitochondrial proteins are synthesized at the cytosolic ribosomes and then imported inside the mitochondria. Concerning the cellular environment, nascent mitochondrial precursor polypeptides may associate with the import machinery while still being synthesized on the ribosome (co-translational import) or may be first released from the ribosome after translation is completed and only then interact with the mitochondria (post-translational import). Most likely, also depending on the individual properties of the preproteins, the *in vivo* situation is represented by a mixture of both processes (Verner, 1993; Mukhopadhyay *et al.*, 2004). As discussed above, in particular Aβ42 showed a strong inhibitory effect in standard post-translational import experiments. After performing import in a co-translational fashion, we found that the mitochondrial import was still inhibited by Aβ peptides although with a lower efficiency than under post-translational conditions. We conclude that the import inhibitory effect of Aβ peptides mainly affects a very initial step of the process when newly synthesized mitochondrial polypeptides are exposed to the cytosolic environment.

Up to date only scarce information is available about direct effects of Aβ peptides on the mitochondrial protein biogenesis process. Using flow cytometry, it was demonstrated that after long-term exposure to Aβ peptides, differentiated PC12 cells exhibited a reduction of newly synthesized mitochondrially-targeted GFP (Sirk *et al.*, 2007). These results are generally in line with our observations, however, due to the long exposure to potentially toxic molecules, these experiments could not distinguish if the import inhibition was a direct or indirect consequence of the presence of Aβ peptides. The immediate inhibitory effect of Aβ peptides on the import reaction in healthy mitochondria, as observed in our experiments, essentially rules out that the inhibition was caused indirectly by a long-term accumulation of functional defects in the affected mitochondria. A single previous study also used isolated mitochondria pretreated for short time with Aβ peptides, but did not detect a deficiency of the mitochondrial import (Hansson Petersen *et al.*, 2008). However, the discrepancy can be explained by an insufficient amount of Aβ peptides used in this study to observe a significant import inhibition. Interestingly, an inhibition of mitochondrial protein biogenesis has been suggested recently as a potential cause for Huntington’s disease (Yano *et al.*, 2014). lt was observed that a mutant form of the protein huntingtin partially inhibited mitochondrial import through a physical association with the TIM23 translocase complex. The concentration of the huntingtin sufficient to obtain inhibition was comparable to the Aβ peptide concentration used in our model.

A recent study proposed that Aβ peptides indirectly interfered with the processing of imported precursor proteins to the mature and active forms (Mossmann *et al.*, 2014), which is an important late step of the mitochondrial import reaction. The authors claimed that an inhibition of PreP (or its yeast homolog Cym1) by Aβ peptides (Alikhani *et al.*, 2011) would result in the accumulation of prepeptides in the mitochondrial matrix that in turn would interfere with the activity of the processing peptidase MPP, required for the maturation of mitochondrial precursor proteins. This is in strong contrast to our study that showed that Aβ peptides acted on an early step of the import reaction. Two observations from our study directly argue against a mitochondrial processing defect caused by Aβ peptides. First, the precursor form visible in import experiments after Aβ peptide inhibition was always sensitive to digestion by external proteases, indicating that the preproteins never crossed the mitochondrial membranes. Second, using two-step import experiments, which separated the binding from the translocation and processing reaction, we observed an inhibitory effect of Aβ peptides only in the first step that is independent of the membrane potential, but not in the second translocation step into the matrix that would also comprise the processing reaction. Although Mossmann et al. found an impaired precursor protein processing activity in presence of Aβ peptides using soluble mitochondrial extracts from yeast as well as in total brain extracts from a murine AD model, the relevance of the claimed processing inhibition for the *in vivo* situation is questionable. Interestingly, they also observed a minor accumulation of precursor polypeptides after cellular expression of Aβ in intact yeast cells and also in brain extracts from AD patients. As a cytosolic accumulation of unprocessed precursor forms is the typical hallmark of a defective overall import process, this observation is strongly consistent with our results of a direct inhibitory effect of Aβ peptides on preprotein import but not processing.

Although a previous experiment indicated a specific and complete import of Aβ peptides into mitochondria (Hansson Petersen *et al.*, 2008), we revisited this question by analyzing the biochemical properties of the interaction of Aβ peptides with isolated and energized mitochondria. Also in our experiments, Aβ42 exhibited some co-sedimentation with mitochondria during the differential centrifugation procedure typically used to re-isolate mitochondria after an import experiment. In addition, Aβ42, but also Aβ40, showed some degree of resistance against added proteases, both observations superficially arguing for a successful import reaction. However, our analysis clearly showed that both Aβ peptides were not taken up by mitochondria because they did not completely satisfy the required criteria of mitochondrial import reaction. Most importantly, the sedimentation behavior and the partial protease resistance of Aβ42 were largely maintained in the absence of mitochondria (mock samples), correlating with its intrinsic tendency to form aggregates. In line with our results are data from the literature showing that both Aβ peptides extracted from AD brains as well as synthetic Aβ peptides spiked into brain homogenates acquired detergent-insolubility and resistance to protease digestion (Soto and Castano, 1996; Xiao *et al.*, 2014). All together these results exclude a complete import of Aβ peptides into mitochondria, but not a peripheral association between Aβ peptides, in particular Aβ42 with the OMM. Our experiments indicated that the presence of mitochondria promoted both aggregation propensity and protease-resistance of Aβ42 (Murphy, 2007; Henry *et al.*, 2015).

Generally, Aβ peptides have an intrinsic tendency to self-assemble into a range of different aggregates also under the conditions that we applied in our mitochondrial import assay (Snyder *et al.*, 1994; Stine *et al.*, 2003; Thal *et al.*, 2015). Utilizing density gradient centrifugation as a method to separate protein aggregates from cell organelles like mitochondria (Sehlin *et al.*, 2012), we observed that a fraction of the Aβ42 peptide added to the experiment directly associated with mitochondria. Interestingly, the presence of precursor proteins changed the behavior of Aβ42 as the amount of mitochondria-associated material decreased while the aggregated forms increased. Additionally, in the presence of Aβ42 a considerable amount of the precursor protein itself was found in the aggregate fraction, indicating the formation of co-aggregates. We propose that this co-aggregation of precursor proteins and Aβ peptides is the main reason for the strong inhibitory effect of mitochondrial protein import. A formation of high molecular weight aggregates and the concomitant reduction of the solubility would significantly reduce their import competence of precursor proteins. Several further observations support this co-aggregation model. Correlated with the much stronger import inhibitory effect of Aβ42 compared to Aβ40, the co-aggregation phenomenon was particularly pronounced in presence of Aβ42. The solubility of the precursor proteins was reduced in presence of Aβ42 as assayed by a centrifugation assay. Together with Aβ42, precursor proteins formed large aggregates that were retarded in a filtration assay. In native PAGE experiments, precursor protein signals were shifted to a high molecular weight complex in the range of 700 kDa that co-purified with Aβ42. Interestingly, recent results showing negative consequences of co-aggregation between cytosolic enzymes and Aβ peptides support this AD-specific pathological mechanism (Itakura *et al.*, 2015). Our work therefore adds an important aspect concerning the deleterious consequences of co-aggregation processes during the etiology of neurodegenerative diseases. Many amyloid diseases involve co-aggregation of different protein species (Penke *et al.*, 2012; Sarell *et al.*, 2013), although the pathological mechanisms are not always entirely clear. It is conceivable that amyloidogenic β-sheet peptides interact with many different endogenous proteins leading to sequestration and functional impairment (Olzscha *et al.*, 2011).

Considering the intracellular space as a crowded environment, Aβ peptides likely undergo multiple, largely non-specific interactions with any protein and lipid components of the cytosol. The import-competent state of mitochondrial preproteins is represented by an incompletely folded conformation that is prone to irregular interactions with Aβ peptides and subsequent aggregation. Already during the onset of the disease at the point at which the concentration of Aβ peptides is increasing, the formation of co-aggregates with newly synthesized mitochondrial precursor polypeptides might progressively interfere with the import process. This would eventually result in a reduction or even loss of mitochondrial enzyme activities, in turn leading to the pleiotropic nature of mitochondrial dysfunction observed in AD patients and respective disease models (Wang *et al.*, 2007; Kaminsky *et al.*, 2015). Hence, the observed strong inhibitory effect on mitochondrial protein import, in particular in case of the pathogenic Aβ42, strongly supports the hypothesis of a direct mitochondrial toxicity of Aβ peptides on mitochondria in AD.

## Material and Methods

### Preparation of Aβ peptides and mitochondrial treatment

The *Escherichia Coli* expressed human recombinant Aβ peptides 1-40 (Ultra Pure HFIP; cat. A-1153-2) and 1-42 (Ultra Pure HFIP; cat. A-1163-2) used in this study were purchased from AJ Roboscreen GmbH (Leipzig, DE). Working solutions of both peptides were prepared as described (Stine *et al.*, 2003). Briefly, the lyophilized peptides were dissolved in 100% 1,1,1,3,3,3-Hexafluoro-2-Propanol (Sigma-Aldrich) and distributed in low-binding micro-centrifuge tubes (VWR, DE). The solvent was allowed to evaporate over night at room temperature and the Aβ peptide aliquots were stored at −80°C. Immediately prior to use, each aliquot was warmed to room temperature followed by a resuspension of the peptide film to a stock of 5 mM in dimethyl sulfoxide (AppliChem GmbH, DE) to remove any preexisting aggregated structures and to provide a homogeneous non-aggregated peptide preparation. After mixing well, the Aβ peptide DMSO stock was freshly diluted with ice-cold distilled water to a final concentration of 100 μM. This dilution was mixed and used immediately. All experiments with Aβ peptides were performed in super-clear tubes (VWR, DE). In some of the experiments, Aβ peptides were precipitated with 72% trichloroacetic acid (TCA) followed by tricine-SDS-PAGE, Western blot and immunodecoration to improve the running behavior of the small peptides.

### Cell culture and isolation of mitochondria

HeLa Cells were cultured in RPMI 1640 medium with 10% heat-inactivated fetal calf serum, 2 mM L-glutamine, 100 units/ml penicillin, and 100 μg/ml streptomycin at 37 °C in a saturated humidity atmosphere containing 5% CO_2_. All the chemicals were bought from Gibco, Life Technologies, DE. The mitochondria were isolated from HeLa cells as described (Becker *et al.*, 2012). Briefly, after harvesting and washing in PBS, cells were incubated for 40 min on ice with HMS-A buffer (0.22 M mannitol, 0.07 M sucrose, 0.02 M HEPES pH 7.4, 1 mM EDTA, 0.2% BSA, 1 mM PMFS). Then, cells were homogenized with a glass/Teflon homogenizer (B. Braun Melsungen AG, DE) followed by differential centrifugation steps to isolated mitochondria. The mitochondria were washed and resuspended in HMS-B buffer (0.22 M mannitol, 0.07 M sucrose, 0.02 M HEPES pH 7.4, 1 mM EDTA, 1 mM PMFS).

### Import of radiolabeled preproteins into isolated mitochondria

The import of radiolabeled precursor proteins was performed essentially as described (Becker *et al.*, 2012). Radiolabeled preproteins were synthesized by *in vitro* transcription/translation using the mMESSAGE mMACHINE transcription kit (Life Technologies, DE) and rabbit reticulocyte lysate (Promega, DE) in presence of [^35^S]-methionine/cysteine (PerkinElmer, DE). For the import reaction, mitochondria were diluted in import buffer (20 mM HEPES-KOH, pH 7.4, 250 mM sucrose, 5 mM magnesium acetate, 80 mM potassium acetate, 5 mM KPi, pH 7.4, 7.5 mM glutamate, 5 mM malate, 1 mM DTT, 2 mM ATP) to a final concentration of 50 μg/100μl. In mock samples, Aβ peptides were incubated under the same buffer conditions but without added mitochondria. Where indicated, mitochondrial membrane potential (Δψ_mit_) was dissipated by adding a mixture of 8 μM antimycin A (Sigma-Aldrich, DE), 0.5 μM valinomycin, and 2 μM oligomycin (Sigma-Aldrich, DE). All the import reactions were performed at 30 °C, stopped by addition of 50 μM valinomycin and placing the samples on ice. Non-imported, protease-accessible mitochondrial proteins were digested by incubation with 100 μg/ml trypsin (Seromed, Biochrom KG, DE) for 30 min on ice and terminated by adding 800 μg/ml of trypsin inhibitor (Sigma-Aldrich, DE) and 1 mM PMFS (Carl Roth, DE). Then, mitochondria were washed in import buffer without substrates. Where indicated, samples were treated with 25 μg/ml proteinase K (PK; Carl Roth, DE) on ice for 30 min before the addition of 1 mM PMSF. After centrifugation for 10 min at 12.000xg and 4 °C, mitochondrial pellets were analyzed by tricine-SDS-PAGE, Western blot, digital autoradiography and immunodecoration.

For two-step import reactions, mitochondrial inner membrane potential Δψ_mit_ was first depleted with 1 μM carbonyl cyanide m-chlorophenyl hydrazone (CCCP). Mitochondria were incubated with radiolabeled preprotein for 30 minutes at 30 °C. After washing, the mitochondria were re-incubated for 30 minutes at 30 °C in energized import buffer supplemented with 2 mg/ml BSA to restore the membrane potential in presence or absence of 3.5 μM Aβ peptides. After re-isolation of mitochondria, imported proteins were separated by tricine-SDS-PAGE and detected by immunodecoration and digital autoradiography.

### BN-PAGE

To analyze mitochondrial protein complexes and Aβ peptide aggregation states under native conditions, samples were analyzed by blue native (BN)-PAGE (Wittig *et al.*, 2006). Isolated mitochondria, as well as co-aggregates containing Aβ peptides and radiolabeled preproteins were solubilized in BN-lysis buffer (20 mM Tris-HCl, pH 7.4, 50 mM NaCl, 10% glycerol, 1 mM EDTA, 1% digitonin, 1 mM PMFS). BN gel loading buffer (100 mM Bis-Tris, pH 7.0, 500 mM ε-amino-n-caproic acid, 5% w/v Coomassie Brilliant Blue G250) was added and the samples were loaded on 5-16.5% BN gels. Native unstained protein standard (Novex, Life Technologies, DE) was used to estimate molecular weights of protein complexes. After running over-night, gels were equilibrated in SDS buffer (1% (w/v) SDS, 0.19 M glycine, 25 mM Tris) and blotted on PDVF membrane (Carl Roth GmbH, DE) followed by immunodecoration and digital autoradiography.

### Sodium carbonate extraction

After incubation of isolated and intact mitochondria with 3.5 μM Aβ peptides, a further incubation in 0.1 M Na_2_CO_3_ solution (pH 11) was performed on ice for 30 min. Then, after withdrawal of a total sample, an ultra-centrifugation step was done in a Beckman TLA-55 at 45000 RPM (123,000 xg) for 40 min at 4 °C. The pellets were resuspended in tricine sample buffer while the supernatants were precipitated with 72% trichloroacetic acid (TCA) followed by tricine-SDS-PAGE, western blot and immunodecoration.

### Sucrose density gradient centrifugation

After incubation with Aβ peptides (35 μM) and/or [^35^S]Su9(70)-DHFR, isolated mitochondria and mock samples were loaded on a continuous sucrose gradient (25-50%) and centrifuged in a Beckman SW41 rotor at 33,000 rpm (135,000 xg) for 1h at 4° C. Then, fractions of 500 μl were collected from the top of each gradient followed by 72% TCA precipitation. Protein pellets were resuspended in tricine loading buffer, separated by tricine-SDS-PAGE and analyzed by Western blot and immunodecoration.

### Membrane potential measurement in isolated mitochondria

Mitochondrial membrane potential (Δψ_mit_) was analyzed by potential-sensitive fluorescent dye tetramethylrhodamine ethyl ester (TMRE) (Molecular Probes, Invitrogen, DE). After incubation with Aβ peptides, isolated mitochondria were resuspended in potential buffer (0.6 M sorbitol, 0.1% BSA, 10 mM MgCl_2_, 20 mM KPi, pH 7.2, 5 mM malate, 10 mM glutamate) and incubated with 1 μM of TMRE for 30 min at 30 °C on ice. After washing away the excess of TMRE, the TMRE fluorescence was measured in a microplate reader (excitation 540 nm, emission 585 nm; Infinite M200 PRO, TECAN, DE).

### Filter retardation assay

To visualize the formation of aggregates and co-aggregates, a modified filter retardation assay (Scherzinger *et al.*, 1997) was used. After incubation of radiolabeled precursor proteins with different amounts of Aβ peptides for 30 min at 30 °C in energized import buffer, samples were filtered directly through cellulose acetate membrane (0.2 μm pore size; GE Healthcare, DE) or nitrocellulose membrane (GE Healthcare, DE) using a dot blot filtration unit (SCIE-PLAS, DE). Proteins retarded on the membranes were analyzed by immunodecoration and digital autoradiography.

### Miscellaneous methods

All the chemicals using in this study were from Carl Roth GmbH or Sigma-Aldrich. Standard techniques were used for tricine-SDS-PAGE, Western blot, and immunodecoration. After performing a tricine-SDS-PAGE, samples were transferred on PVDF membrane (Carl Roth GmbH) followed by blocking in TBS/Tween (0.9% NaCl, 10 mM Tris/HCl pH 7.4, 0.25% Tween 20) with 5% milk and immunodecoration with andibodies appropriately diluted in TBS/Tween. Signal detection was performed by enhanced chemiluminence (SERVA Light Eos Ultra, Serva, DE). Used antibodies were: Aβ 6E10 (Covance SIG-39320); Tim23 (BD Bioscience 611222), Tom 20 (Santa Cruz SC-11415), Tom 40 (Santa Cruz SC-11414), SMAC (Santa Cruz SC-22766), MPP (Sigma-Aldrich HPA021648), Complex-I (Invitrogen 459100), Complex-II (Invitrogen 459200), Complex III (Santa Cruz SC-23986), Complex-IV (Cell Signaling 3E11), F_1_β (Invitrogen A21351), Rabbit IgG-Peroxidase (Sigma Aldrich A6154) and Mouse IgG-Peroxidase (Sigma Aldrich A4416). Digital autoradiography was performed using a FLA5100 phosphorimaging system (Fujifilm, DE). Quantitative analysis was done by ImageJ 64 (NIH, USA) and GraphPad Prism 6.0 (GraphPad Software, Inc, USA).

## Acknowledgements

We are grateful to U. Gerken for expert technical support and Prof. J. Walter from University of Bonn (Germany) for his critical comments. This work was supported by a fellowship from European Commission Framework Programs (Marie Curie Action 7th Framework Program - International Incoming Fellowship; to GC) and by a research grant from the Deutsche Forschungsgemeinschaft (VO657/5-2; to WV).

## References

Agostinho, P., Pliassova, A., Oliveira, C.R., and Cunha, R.A. (2015). Localization and trafficking of amyloid-beta protein precursor and secretases: Impact on Alzheimer’s disease. J. Alzheimers Dis. 45, 329–347.

Alikhani, N., Guo, L., Yan, S., Du, H., Pinho, C.M., Chen, J.X., Glaser, E., and Yan, S.S. (2011). Decreased proteolytic activity of the mitochondrial amyloid-beta degrading enzyme, PreP peptidasome, in Alzheimer’s disease brain mitochondria. J. Alzheimers Dis. 27, 75–87.

Becker, D., Richter, J., Tocilescu, M.A., Przedborski, S., and Voos, W. (2012). Pink1 kinase and its membrane potential (Deltapsi)-dependent cleavage product both localize to outer mitochondrial membrane by unique targeting mode. J. Biol. Chem. 287, 22969–22987.

Bitan, G., Kirkitadze, M.D., Lomakin, A., Vollers, S.S., Benedek, G.B., and Teplow, D.B. (2003). Amyloid beta -protein (Abeta) assembly: Abeta 40 and Abeta 42 oligomerize through distinct pathways. Proc. Natl. Acad. Sci. USA 100, 330–335.

Chacinska, A., Koehler, C.M., Milenkovic, D., Lithgow, T., and Pfanner, N. (2009). Importing mitochondrial proteins: machineries and mechanisms. Cell 138, 628–644.

Eckman, C.B., and Eckman, E.A. (2007). An update on the amyloid hypothesis. Neurol. Clin. 25, 669–682, vi.

Endo, T., and Kohda, D. (2002). Functions of outer membrane receptors in mitochondrial protein import. Biochim. Biophys. Acta 1592, 3–14.

Fox, T.D. (2012). Mitochondrial protein synthesis, import, and assembly. Genetics 192, 1203–1234.

Gouras, G.K., Tampellini, D., Takahashi, R.H., and Capetillo-Zarate, E. (2010). Intraneuronal beta-amyloid accumulation and synapse pathology in Alzheimer’s disease. Acta Neuropathol. 119, 523–541.

Haass, C., and Selkoe, D.J. (2007). Soluble protein oligomers in neurodegeneration: lessons from the Alzheimer’s amyloid beta-peptide. Nat. Rev. Mol. Cell Biol. 8, 101–112.

Hansson Petersen, C.A., Alikhani, N., Behbahani, H., Wiehager, B., Pavlov, P.F., Alafuzoff, I., Leinonen, V., Ito, A., Winblad, B., Glaser, E., and Ankarcrona, M. (2008). The amyloid beta-peptide is imported into mitochondria via the TOM import machinery and localized to mitochondrial cristae. Proc. Natl. Acad. Sci. USA 105, 13145–13150.

Hardy, J., and Selkoe, D.J. (2002). The amyloid hypothesis of Alzheimer’s disease: progress and problems on the road to therapeutics. Science 297, 353–356.

Hardy, J.A., and Higgins, G.A. (1992). Alzheimer’s disease: the amyloid cascade hypothesis. Science 256, 184–185.

Head, E., and Lott, I.T. (2004). Down syndrome and beta-amyloid deposition. Curr. Opin. Neurol. 17, 95–100.

Heiser, V., Scherzinger, E., Boeddrich, A., Nordhoff, E., Lurz, R., Schugardt, N., Lehrach, H., and Wanker, E.E. (2000). Inhibition of huntingtin fibrillogenesis by specific antibodies and small molecules: implications for Huntington’s disease therapy. Proc. Natl. Acad. Sci. USA 97, 6739–6744.

Henry, S., Vignaud, H., Bobo, C., Decossas, M., Lambert, O., Harte, E., Alves, I.D., Cullin, C., and Lecomte, S. (2015). Interaction of Abeta(1–42) amyloids with lipids promotes “off-pathway” oligomerization and membrane damage. Biomacromolecules 16, 944–950.

Itakura, M., Nakajima, H., Kubo, T., Semi, Y., Kume, S., Higashida, S., Kaneshige, A., Kuwamura, M., Harada, N., Kita, A., Azuma, Y.T., Yamaji, R., Inui, T., and Takeuchi, T. (2015). Glyceraldehyde-3-phosphate dehydrogenase aggregates accelerate amyloid-beta amyloidogenesis in Alzheimer disease. J. Biol. Chem. 290, 26072–26087.

Jarrett, J.T., Berger, E.P., and Lansbury, P.T., Jr. (1993). The C-terminus of the beta protein is critical in amyloidogenesis. Ann. NY Acad. Sci. 695, 144–148.

Kaminsky, Y.G., Tikhonova, L.A., and Kosenko, E.A. (2015). Critical analysis of Alzheimer’s amyloid-beta toxicity to mitochondria. Front. Biosci. 20, 173–197.

Kummer, M.P., and Heneka, M.T. (2014). Truncated and modified amyloid-beta species. Alzheimers Res. Ther. 6, 28.

LaFerla, F.M., Green, K.N., and Oddo, S. (2007). Intracellular amyloid-beta in Alzheimer’s disease. Nat. Rev. Neurosci. 8, 499–509.

Lustbader, J.W., Cirilli, M., Lin, C., Xu, H.W., Takuma, K., Wang, N., Caspersen, C., Chen, X., Pollak, S., Chaney, M., Trinchese, F., Liu, S., Gunn-Moore, F., Lue, L.F., Walker, D.G., Kuppusamy, P., Zewier, Z.L., Arancio, O., Stern, D., Yan, S.S., and Wu, H. (2004). ABAD directly links Abeta to mitochondrial toxicity in Alzheimer’s disease. Science 304, 448–452.

Mattson, M.P., Gleichmann, M., and Cheng, A. (2008). Mitochondria in neuroplasticity and neurological disorders. Neuron 60, 748–766.

Mawuenyega, K.G., Sigurdson, W., Ovod, V., Munsell, L., Kasten, T., Morris, J.C., Yarasheski, K.E., and Bateman, R.J. (2010). Decreased clearance of CNS beta-amyloid in Alzheimer’s disease. Science 330, 1774.

Mossmann, D., Vogtle, F.N., Taskin, A.A., Teixeira, P.F., Ring, J., Burkhart, J.M., Burger, N., Pinho, C.M., Tadic, J., Loreth, D., Graff, C., Metzger, F., Sickmann, A., Kretz, O., Wiedemann, N., Zahedi, R.P., Madeo, F., Glaser, E., and Meisinger, C. (2014). Amyloid-beta peptide induces mitochondrial dysfunction by inhibition of preprotein maturation. Cell Metab. 20, 662–669.

Mukhopadhyay, A., Ni, L., and Weiner, H. (2004). A co-translational model to explain the in vivo import of proteins into HeLa cell mitochondria. Biochem. J. 382, 385–392.

Murphy, M.P., and LeVine, H., 3rd. (2010). Alzheimer’s disease and the amyloid-beta peptide. J. Alzheimers Dis. 19, 311–323.

Murphy, R.M. (2007). Kinetics of amyloid formation and membrane interaction with amyloidogenic proteins. Biochim Biophys Acta 1768, 1923–1934.

Musiek, E.S., and Holtzman, D.M. (2015). Three dimensions of the amyloid hypothesis: time, space and ‘wingmen’. Nat. Neurosci. 18, 800–806.

Olzscha, H., Schermann, S.M., Woerner, A.C., Pinkert, S., Hecht, M.H., Tartaglia, G.G., Vendruscolo, M., Hayer-Hartl, M., Hartl, F.U., and Vabulas, R.M. (2011). Amyloid-like aggregates sequester numerous metastable proteins with essential cellular functions. Cell 144, 67–78.

Pagani, L., and Eckert, A. (2011). Amyloid-Beta interaction with mitochondria. Int. J. Alzheimers Dis. 2011, 925050.

Penke, B., Toth, A.M., Foldi, I., Szucs, M., and Janaky, T. (2012). Intraneuronal beta-amyloid and its interactions with proteins and subcellular organelles. Electrophoresis 33, 3608–3616.

Piaceri, I., Rinnoci, V., Bagnoli, S., Failli, Y., and Sorbi, S. (2012). Mitochondria and Alzheimer’s disease. J. Neurol. Sci. 322, 31–34.

Roher, A.E., Esh, C.L., Kokjohn, T.A., Castano, E.M., Van Vickle, G.D., Kalback, W.M., Patton, R.L., Luehrs, D.C., Daugs, I.D., Kuo, Y.M., Emmerling, M.R., Soares, H., Quinn, J.F., Kaye, J., Connor, D.J., Silverberg, N.B., Adler, C.H., Seward, J.D., Beach, T.G., and Sabbagh, M.N. (2009). Amyloid beta peptides in human plasma and tissues and their significance for Alzheimer’s disease. Alzheimers Dement 5, 18–29.

Ryan, M.T., Voos, W., and Pfanner, N. (2001). Assaying protein import into mitochondria. Methods Cell Biol. 65, 189–215.

Sannerud, R., and Annaert, W. (2009). Trafficking, a key player in regulated intramembrane proteolysis. Semin Cell Dev Biol 20, 183–190.

Sarell, C.J., Stockley, P.G., and Radford, S.E. (2013). Assessing the causes and consequences of co-polymerization in amyloid formation. Prion 7, 359–368.

Scherzinger, E., Lurz, R., Turmaine, M., Mangiarini, L., Hollenbach, B., Hasenbank, R., Bates, G.P., Davies, S.W., Lehrach, H., and Wanker, E.E. (1997). Huntingtin-encoded polyglutamine expansions form amyloid-like protein aggregates in vitro and in vivo. Cell 90, 549–558.

Sehlin, D., Englund, H., Simu, B., Karlsson, M., Ingelsson, M., Nikolajeff, F., Lannfelt, L., and Pettersson, F.E. (2012). Large aggregates are the major soluble Abeta species in AD brain fractionated with density gradient ultracentrifugation. PLoS One 7, e32014.

Selfridge, J.E., E, L., Lu, J., and Swerdlow, R.H. (2013). Role of mitochondrial homeostasis and dynamics in Alzheimer’s disease. Neurobiol. Dis. 51, 3–12.

Sirk, D., Zhu, Z., Wadia, J.S., Shulyakova, N., Phan, N., Fong, J., and Mills, L.R. (2007). Chronic exposure to sub-lethal beta-amyloid (Abeta) inhibits the import of nuclear-encoded proteins to mitochondria in differentiated PC12 cells. J. Neurochem. 103, 1989–2003.

Snyder, S.W., Ladror, U.S., Wade, W.S., Wang, G.T., Barrett, L.W., Matayoshi, E.D., Huffaker, H.J., Krafft, G.A., and Holzman, T.F. (1994). Amyloid-beta aggregation: selective inhibition of aggregation in mixtures of amyloid with different chain lengths. Biophys. J. 67, 1216–1228.

Soto, C., and Castano, E.M. (1996). The conformation of Alzheimer’s beta peptide determines the rate of amyloid formation and its resistance to proteolysis. Biochem. J. 314 (Pt 2), 701–707.

Stine, W.B., Jr., Dahlgren, K.N., Krafft, G.A., and LaDu, M.J. (2003). In vitro characterization of conditions for amyloid-beta peptide oligomerization and fibrillogenesis. J. Biol. Chem. 278, 11612–11622.

Swerdlow, R.H., Burns, J.M., and Khan, S.M. (2014). The Alzheimer’s disease mitochondrial cascade hypothesis: progress and perspectives. Biochim. Biophys. Acta. 1842, 1219–1231.

Thal, D.R., Walter, J., Saido, T.C., and Fandrich, M. (2015). Neuropathology and biochemistry of Abeta and its aggregates in Alzheimer’s disease. Acta Neuropathol. 129, 167–182.

Truscott, K.N., Wiedemann, N., Rehling, P., Muller, H., Meisinger, C., Pfanner, N., and Guiard, B. (2002). Mitochondrial import of the ADP/ATP carrier: the essential TIM complex of the intermembrane space is required for precursor release from the TOM complex. Mol. Cell. Biol. 22, 7780–7789.

Verner, K. (1993). Co-translational protein import into mitochondria: an alternative view. Trends Biochem. Sci. 18, 366–371.

Wang, X., Su, B., Perry, G., Smith, M.A., and Zhu, X. (2007). Insights into amyloid-beta-induced mitochondrial dysfunction in Alzheimer disease. Free Radic. Biol. Med. 43, 1569–1573.

Weller, R.O., Massey, A., Kuo, Y.M., and Roher, A.E. (2000). Cerebral amyloid angiopathy: accumulation of A beta in interstitial fluid drainage pathways in Alzheimer’s disease. Ann. NY Acad. Sci. 903, 110–117.

Wirths, O., and Bayer, T.A. (2012). Intraneuronal Abeta accumulation and neurodegeneration: lessons from transgenic models. Life Sci 91, 1148–1152.

Wirths, O., Multhaup, G., and Bayer, T.A. (2004). A modified beta-amyloid hypothesis: intraneuronal accumulation of the beta-amyloid peptide--the first step of a fatal cascade. J. Neurochem. 91, 513–520.

Wittig, I., Braun, H.P., and Schägger, H. (2006). Blue native PAGE. Nat. Protoc. 1, 418–428.

Xiao, X., Yuan, J., Qing, L., Cali, I., Mikol, J., Delisle, M.B., Uro-Coste, E., Zeng, L., Abouelsaad, M., Gazgalis, D., Martinez, M.C., Wang, G.X., Brown, P., Ironside, J.W., Gambetti, P., Kong, Q., and Zou, W.Q. (2014). Comparative Study of Prions in Iatrogenic and Sporadic Creutzfeldt-Jakob Disease. J. Clin. Cell. Immunol. 5.

Yano, H., Baranov, S.V., Baranova, O.V., Kim, J., Pan, Y., Yablonska, S., Carlisle, D.L., Ferrante, R.J., Kim, A.H., and Friedlander, R.M. (2014). Inhibition of mitochondrial protein import by mutant huntingtin. Nat. Neurosci. 17, 822–831.

Zhang, Y.W., Thompson, R., Zhang, H., and Xu, H. (2011). APP processing in Alzheimer’s disease. Mol. Brain 4, 3.

